# Fat1 regulates astrocyte maturation and angiogenesis in the retina

**DOI:** 10.1101/2020.04.23.057141

**Authors:** Françoise Helmbacher

**Affiliations:** Aix Marseille Univ, CNRS, IBDM UMR 7288, Parc Scientifique de Luminy, Case 907, 13288 Marseille, France

## Abstract

Angiogenesis is a stepwise process leading to blood vessel formation. In the vertebrate retina, endothelial cells are guided by astrocytes migrating along the inner surface, and the two processes are coupled by a tightly regulated cross-talk between the two cell types. Here, we investigated how the FAT1 Cadherin, a regulator of tissue morphogenesis governing tissue cross-talks, influences retinal vascular development. Through late-onset inactivation in the neural lineage in mice, we bypassed an early contribution of *Fat1* to eye development, and assessed its requirement for postnatal retina angiogenesis. We found that neural *Fat1* expression, by controlling the polarity of astrocyte progenitor migration, regulates astrocyte maturation. By interfering with astrocyte migration and maturation, neural *Fat1* deletion deregulates the astrocyte/endothelial cell coupling, and delays retinal angiogenesis. Mice with neural-*Fat1* ablation exhibit persistent abnormalities of the retinal vascular architecture, such as an increased vascular density in deep layers. Altogether, this study identifies *Fat1* as a regulator of neurovascular communication, essential for retinal vascular development and integrity.

## Introduction

The vascular system is dedicated to the distribution of nutrients and oxygen to all organs, and serves as a vector for hormonal control and immune surveillance. Assembly of the vascular system involves a process called angiogenesis, by which endothelial cells coalesce to form vessels, which are subsequently remodeled so as to adapt blood flow to tissue scale. Whereas the cellular and molecular mechanisms underlying these processes are common to many organs (Ruhrberg and Bautch, 2013), some vascular properties, such as the capacity of exchange and filtration, are adjusted in an organ-specific manner, and such adaptations involve tissue-specific cross-talks between endothelial cells and cellular players. An emblematic example of organ-specific adaptation is the tight barrier function by which central nervous system vasculature restricts cellular permeability and immune passage, and this specificity is driven by numerous exchanges of signals between neural and endothelial cells (Paredes et al., 2018).

The retina represents an excellent system for the study of these neurovascular interactions. In the retina, angiogenesis starts after birth, involving first a phase (between P0 and P7) of horizontal vascular growth in the plane of the inner-most retinal layer (sharing the space with axons of retinal ganglion cell (RGC) neurons, as they migrate towards the optic nerve). This process is followed by a phase of perpendicular growth of vessels towards deeper retinal layers, (between P7 and P21), resulting in the successive formation of two additional vascular layers, called inner and outer plexiform layers, respectively (Paredes et al., 2018; Ruhrberg and Bautch, 2013). During the phase of horizontal vascular growth, endothelial cells (ECs) are preceded by migrating astrocyte progenitor cells (APCs), which enter the retina through the optic nerve (Watanabe and Raff, 1988), and spread radially on the inner retinal surface, reaching the edges of the retina in a few days, ahead of EC migration (Tao and Zhang, 2014). Astrocyte motility is ensured by PDGF, secreted by the underlying RGC neurons, and signaling via PDGFRα (Fruttiger et al., 1996). APCs subsequently undergo partial differentiation, and turn into immature astrocytes, which serve as substrate and guidance cue for the subsequent migration of endothelial cells. Ahead of the EC migration front, immature astrocytes secrete vascular endothelial growth factor (VEGF) in response to hypoxia (Stone et al., 1995). VEGF acts as a chemo-attractant for EC migration, signaling via its cognate receptors VEGFRs (Gerhardt et al., 2003).

As ECs migrate radially on the astrocytic layer, allowing the vascular network to expand, the rise in oxygen level resulting from blood perfusion in newly formed vessels locally promotes the conversion of immature astrocytes into mature astrocytes, characterized by increased expression of the astrocyte marker glial fibrillary acidic protein (GFAP) and reduced VEGF expression levels proximal to the vascular front (West et al., 2005). This oxygen-mediated silencing of VEGF expression in astrocytes in contact with ECs (proximal to the vascular front), is opposed to the high VEGF levels induced by hypoxia in the EC-free distal region ahead of the vascular front (Duan et al., 2017; Duan et al., 2014; Rattner et al., 2019; Stone et al., 1995). This dual regulation establishes a gradient of VEGF (high in immature astrocytes, low in mature astrocytes), determining the outward direction polarity of EC migration (West et al., 2005). Because of this mutual dependency, molecular alterations that interfere with astrocyte migration also impact on the progression and angiogenesis in the postnatal retina (Duan et al., 2017; Duan et al., 2014; Kautzman et al., 2018; Lavina et al., 2018; Tao and Zhang, 2016), whereas manipulations interfering with EC migration also result in defective astrocyte maturation (Morita et al., 2017). Aside from astrocytes, other neuronal cell types and signaling systems also contribute to modulate angiogenesis in the nervous system (Paredes et al., 2018). Neurons participate in several ways, either by sending pro-angiogenic factors, such as VEGF (Himmels et al., 2017; James et al., 2009), indirect contributors such as PDGFA, which acts on astrocytes (Fruttiger et al., 1996; Gerhardt et al., 2003; Tao and Zhang, 2016), or negative regulators of angiogenesis such as the decoy VEGF receptor sFlt1 and Sema3E (Himmels et al., 2017; Kim et al., 2011; Zygmunt et al., 2011). Thus, a complex interplay of pro- and anti-angiogenic factors produced by neural cells is key to orchestrate astrocyte/endothelial cell coupling, hence influencing the timing and dynamics of angiogenesis.

We previously reported that constitutive deletion of the *Fat1*-Cadherin gene, aside from its impact on neuromuscular morphogenesis (Caruso et al., 2013; Helmbacher, 2018), causes abnormalities of the retinal vasculature (Caruso et al., 2013), thus identifying *Fat1* as a putative novel player in vascular biology. However, the exact role played by *Fat1* had not been elucidated. Fat-like cadherins are known regulators of tissue-morphogenesis, capable of modulating cell polarity, cell migration, cell growth, and cell fate decisions (Blair and McNeill, 2018; Horne-Badovinac, 2017; Sadeqzadeh et al., 2014). Their binding to Dachsous-like cadherins triggers bidirectional signaling cascades impinging on the planar cell polarity (PCP) and Hippo/YAP pathways, thereby influencing, both cell- and non-cell-autonomously, cytoskeletal dynamics and cell fate decisions (Blair and McNeill, 2018). Both the Hippo/YAP (Kim et al., 2017; Nakajima et al., 2017; Sakabe et al., 2017; Wang et al., 2017) and the PCP (Cirone et al., 2008) pathways are known modulators of angiogenesis. *Fat1* and *Yap* are also both involved in eye morphogenesis and their inactivation similarly causes a congenital malformation called coloboma (Lahrouchi et al., 2019; Williamson et al., 2014). A particularly well studied activity of Fat-Dachsous signaling is the coordination of cell polarity during events of polarized migration (Horne-Badovinac, 2017). We recently illustrated how *Fat1* coordinates migration and differentiation of myogenic progenitors in the context of a subcutaneous muscle exhibiting characteristics of planar polarized migration, through complementary activities in several cell types (Helmbacher, 2018). Postnatal angiogenesis, with two events of polarized migration coupled with each-other, and known involvements of the classical FAT signaling effectors, represents another logical place where *Fat1* may coordinate morphogenesis.

Here, we explored how *Fat1* influences retinal vascular development in mice. During postnatal angiogenesis, *Fat1* expression is predominantly expressed in astrocytes and Müller glia. Its deletion in the neural lineage was achieved with a late-onset CRE driver to bypass its early morphogenetic activity, thereby rescuing the coloboma found in constitutive knockouts. This led to a delay in the rate of endothelial cell horizontal migration, associated with a delay in astrocyte maturation, manifested by lowered levels of GFAP and PDGFRα, and with altered astrocyte migration polarity at the leading edge. Thus, *Fat1* activity in the neural lineage is required for astrocyte migration polarity and subsequent maturation, and its deletion interferes with the astrocyte/endothelial cell cross-talk, thus delaying retinal angiogenesis. Abnormalities of the vascular architecture persisted at adult stages, making adult mice with neural-specific *Fat1* ablation a new model of neuro-vascular pathology.

## Results

### Constitutive Fat1 disruption impairs eye cup morphogenesis and optic fissure closure

The present study aimed to investigate how *Fat1* gene activity in the postnatal neural lineage contributes to neurovascular cross-talk in the retina. However, *Fat1* disruption is known to affect optic cup development, resulting in morphological abnormalities of the eye. Such defects vary in severity and penetrance, ranging from microphthalmic eyes with high numbers of apoptotic cells and altered lens morphogenesis, to normally shaped eyes exhibiting vascular abnormality in adults (Caruso et al., 2013; Ciani et al., 2003; Lahrouchi et al., 2019; Sugiyama et al., 2015). Such a large range of severity implies not only that *Fat1* is required at distinct developmental stages, but also that the phenotypic penetrance of several knockout alleles is incomplete for alterations of each of these functions. When assessing eye and retinal morphology from E12.-5 to E14.5 in *Fat1*-deficient embryos, we observed a range of phenotypes recapitulating known phenotypes, including ventral coloboma (Fig. 1) (Caruso et al., 2013; Ciani et al., 2003; Lahrouchi et al., 2019; Sugiyama et al., 2015).

**Fig. 1.**
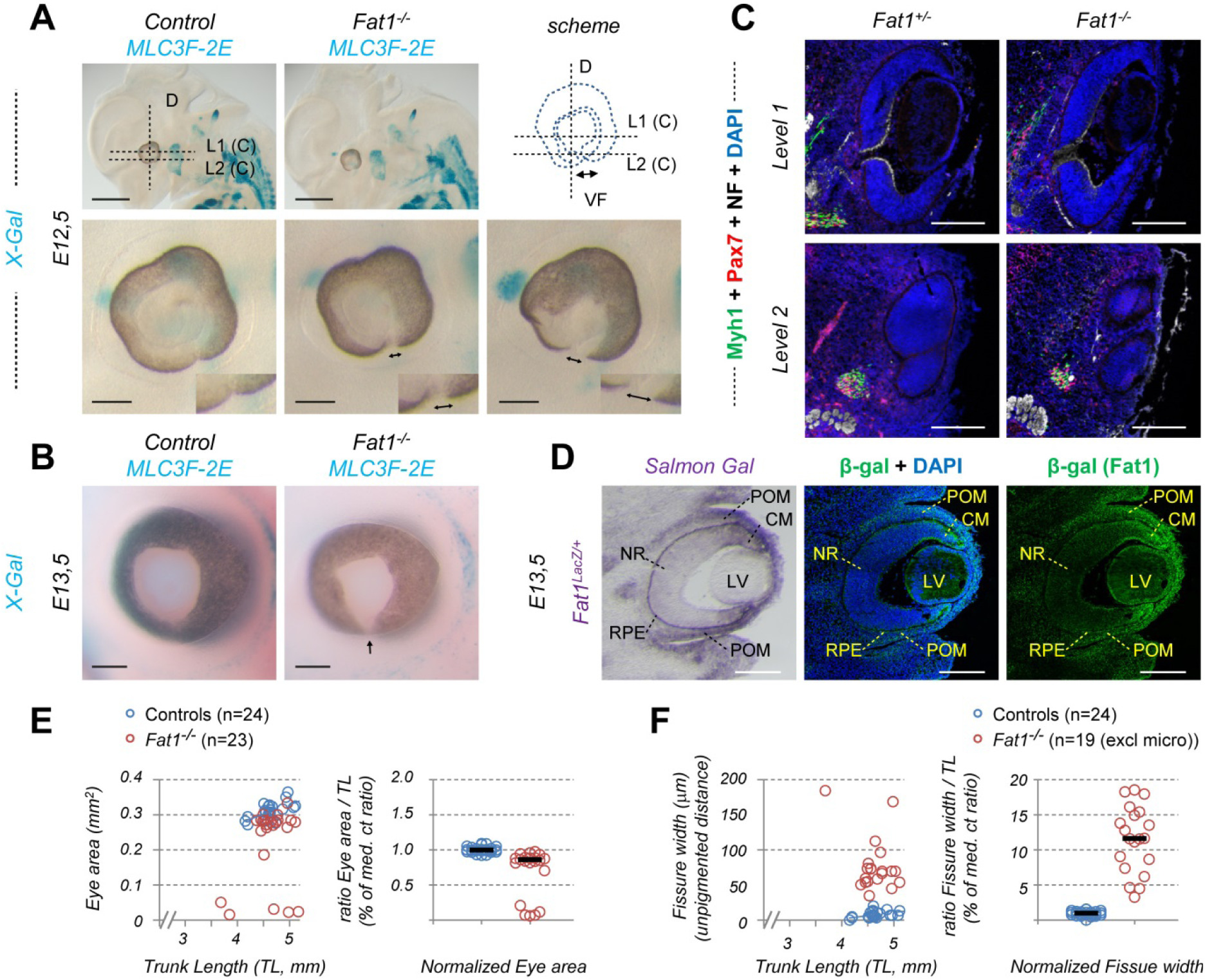
Fat1 disruption causes a ventral Coloboma phenotype. (A, B) Eye morphology was analyzed on whole mount control or *Fat1*^*-/-*^ embryos at E12.5 (A) and E13.5 (B), in which muscles are stained with X-gal owing to the *Mlc3F-2E* transgene. (C) Immunohistochemistry analysis of the retina in wild type and *Fat1*^*-/-*^ embryos E12.5 on successive transverse sections (horizontal with respect to the eye) at two levels of the retina (indicated with horizontal dotted lines labelled as C1 and C2 in (A)), with DAPI nuclear staining (blue), anti-neurofilament (white), anti-Pax7 (red) and anti-Myh1 (green). (D) *Fat1* expression is visualized by Salmon gal staining (left panel), or by anti-β-galactosidase (middle/right panel) immunostaining on neighboring serial coronal sections of a *Fat1*^*LacZ/*+^ E13.5 embryo (angle of sections indicated with a vertical dotted line labelled as D in panel (A)). High levels of *Fat1* expression are detected in periocular mesenchyme (POM), with a ventral hotspot, whereas expression in the neural retina is mostly confined to the ciliary margins (of both the NR and RPE), and to the lens. (E, F) Quantifications of the eye area (EA, E), and the ventral fissure width (VFW, F), relative to the trunk length (TL, proportional to embryo size) are plotted either as raw data (Left plots), or normalized ratios (EA/TL or VFW/TL, relative to median ratio of controls, right plots). CM: ciliary margins; l: lens; NR: neural retina; POM: peri-ocular mesenchyme; RPE: retinal pigmented epithelium; vPOM: ventral periocular mesenchyme. ONL: photoreceptor outer nuclear layer; OPL: outer-plexiform layer; INL: inner nuclear layer. Scale bar: (A): top 1mm; bottom 200μm; (B, C, D): 200μm.

Eye cup morphogenesis starts in the embryo with the emergence of the optic vesicles (OV), connected to the neural tube by structures called optic stalks (Fuhrmann, 2010; Mui et al., 2005). As the OVs grow, their juxtaposition with the head ectoderm triggers a complex rearrangement of the neural ectoderm, which invaginates to form a bi-layered structure, the optic cup (OC) (Tao and Zhang, 2014). The optic cup is constituted of an inner layer, forming the neural retina (NR), and of an outer layer corresponding to the retinal pigmented epithelium (RPE) (Fuhrmann, 2010). Because the optic cup invagination occurs asymmetrically with respect to the optic vesicle, it remains transiently open on its ventral side, the resulting opening being called the optic fissure (OF) (Chang et al., 2006; Fuhrmann, 2010; Morcillo et al., 2006). The OF is later subject to an active morphogenetic process called ventral closure, during which the ventral edges of the optic cup align and fuse, ensuring continuity of the retina on the ventral side (Chang et al., 2006). When the process of ventral closure is defective, the optic cup remains open, causing a congenital abnormality called coloboma, which represents a major cause of childhood blindness (Bovolenta and Martinez-Morales, 2018; Chang et al., 2006).

At E12.5, around the stage of completion of optic fissure closure, control embryos exhibited a nearly closed eye cup, with the pigmented ventral edges closely apposed against each other (Fig. 1A). At that stage, a small proportion of *Fat1*^*-/-*^ embryos (6/23) exhibited severe reduction in the size of the optic cup (microphthalmia), whereas in the majority of embryos (17/23) the eye size (area relative to trunk length) was only modestly reduced (Fig. 1A,E). In these *Fat1*^*-/-*^ embryos without microphthalmic eyes, the optic fissure failed to close, resulting in a coloboma characterized by an increased ventral groove width (Fig. 1A,F), also distinguishable on posterior coronal sections (Fig. 1C). Whereas ventral edges are nearly fused in control embryos, a large grove separates the ventral edges of *Fat1*^*-/-*^ optic cups, with mesenchymal tissue abnormally located in the groove (Fig. 1C). At E14.5, whereas all control eyes exhibited a perfectly continuous ventral retina, *Fat1*^*-/-*^ embryos still presented a distinguishable coloboma (Fig. 1B), indicating that the phenotype observed at E12.5 does not correspond to a delay in the closure, but rather a failure to do so. Thus, *Fat1* activity at early stages of eye morphogenesis is required for optic fissure closure. Such a role was recently shown to be evolutionarily conserved, as *Fat1* loss-of-function mutations lead to coloboma not only in mouse, but also in zebrafish and human, thus adding *FAT1* to the list of known regulators of OF closure in which pathogenic variants are associated with human coloboma (Lahrouchi et al., 2019).

### Delayed Fat1 ablation in the neural lineage bypasses early eye morphogenesis defects

The presence of phenotypes resulting from impaired eye morphogenesis makes the constitutive knockout unsuitable for studying potential *Fat1* involvement at later stages of eye development. Thus, assessing its potential function in cells of the neural lineage implies deleting *Fat1* only after the completion of eye development processes. *Fat1* expression at these stages is not restricted to one single tissue type, but is predominant in the mesenchyme just below the eye cup, in the ciliary margin of the neural retina, and in the RPE (Fig. 1D), as seen by following expression of a *Fat1*^*LacZ*^ allele in embryo sections. Each of these components may contribute significantly to eye growth and optic fissure closure, raising the possibility that retina-specific deletion at an early stage could potentially be detrimental for eye morphogenesis. Furthermore, a recent study linking FAT1 mutations to coloboma attributes this phenotype to *Fat1* activity in the RPE (Lahrouchi et al., 2019). To circumvent this, we used the *Nestin-cre* line, which allows neural-specific CRE-mediated excision, but with a late onset in certain regions resulting in delayed recombination, compared to most other neuronal areas (Lamballe et al., 2011; Tronche et al., 1999). We previously showed that *Nestin-cre*-driven deletion of the HGF receptor gene *Met* in spinal motor neurons, only achieved complete *Met* excision at E13.5 (Lamballe et al., 2011). This enabled preservation of early *Met* function in motor neurons (MNs), so as to address *Met* functions at later stages of MN development (Lamballe et al., 2011). Irrespective of the potential tissue-specificity underlying *Fat1* activity as a regulator of eye morphogenesis, late *Nestin-cre* activity in the retina enables ablation of neural *Fat1* expression late enough to preserve its putative activities during early morphogenesis, at the time of optic fissure closure.

To determine if the timing of *Nestin-cre*-mediated activity in the developing optic cup was compatible with such an approach, we evaluated recombination efficiency with a *Rosa26*-based recombination reporter (*R26-YFP* (Srinivas et al., 2001)), in which YFP expression is conditioned by the CRE-mediated deletion of a Lox-STOP-Lox cassette. Once recombined, YFP expression is permanently driven by the ubiquitous R26 locus, thus allowing tracing of the cells in which CRE was active. We analyzed the pattern of YFP expression achieved in *Nestin-cre*; *R26*^*YFP/*+^ retinas at different stages. At E13.5, one day after completion of the morphogenetic events governing optic fissure fusion, whereas whole embryo visualization of YFP expression outlined the whole central nervous system (CNS), surrounded by otherwise non-recombined cells, illustrating the neural-specificity of *Nestin-cre* activity, little fluorescence could be detected in the eye (Fig. 2A). To further characterize the extent of CRE-mediated excision in the eye, we next performed YFP IHC on embryo sections at E13.5 and E16.5 (Fig. 2A). As expected we detected robust YFP expression in a large fraction of the CNS, and in almost all cells of the proliferative epithelium of the lens vesicle (Fig. 2A, middle panels). In contrast, recombination activity in the E13.5 optic cup itself was limited to a few cells in the ventral RPE, and completely absent in the neural retina at that stage, with no YFP+ cells detected (Fig. 2A, top right panel). These results confirm that at E13.5, when eye morphogenesis is complete, *Nestin-cre*-driven recombination in the eye is restricted to the lens epithelium and ventral RPE, but absent from the neural retina, contrasting with the CNS, where robust recombination has already been achieved. *Nestin-cre* activity only started to be detectable in the neural retina at E16.5, with some YFP^+^ cells present in several retinal layers encompassing glia and neurons (Fig. 2A, bottom panels), indicating that CRE-mediated deletion was ongoing at that stage, while recombination was already near complete in the lens. Finally, in adult retinas, the extent of *Nestin-cre*-mediated deletion reached 100% efficiency in Müller Glia cells, with a complete overlap between YFP and Glutamine Synthetase (GS, *Glul* gene) signals, and in astrocytes (labeled with GFAP antibodies) (Fig. 2B).

**Fig. 2.**
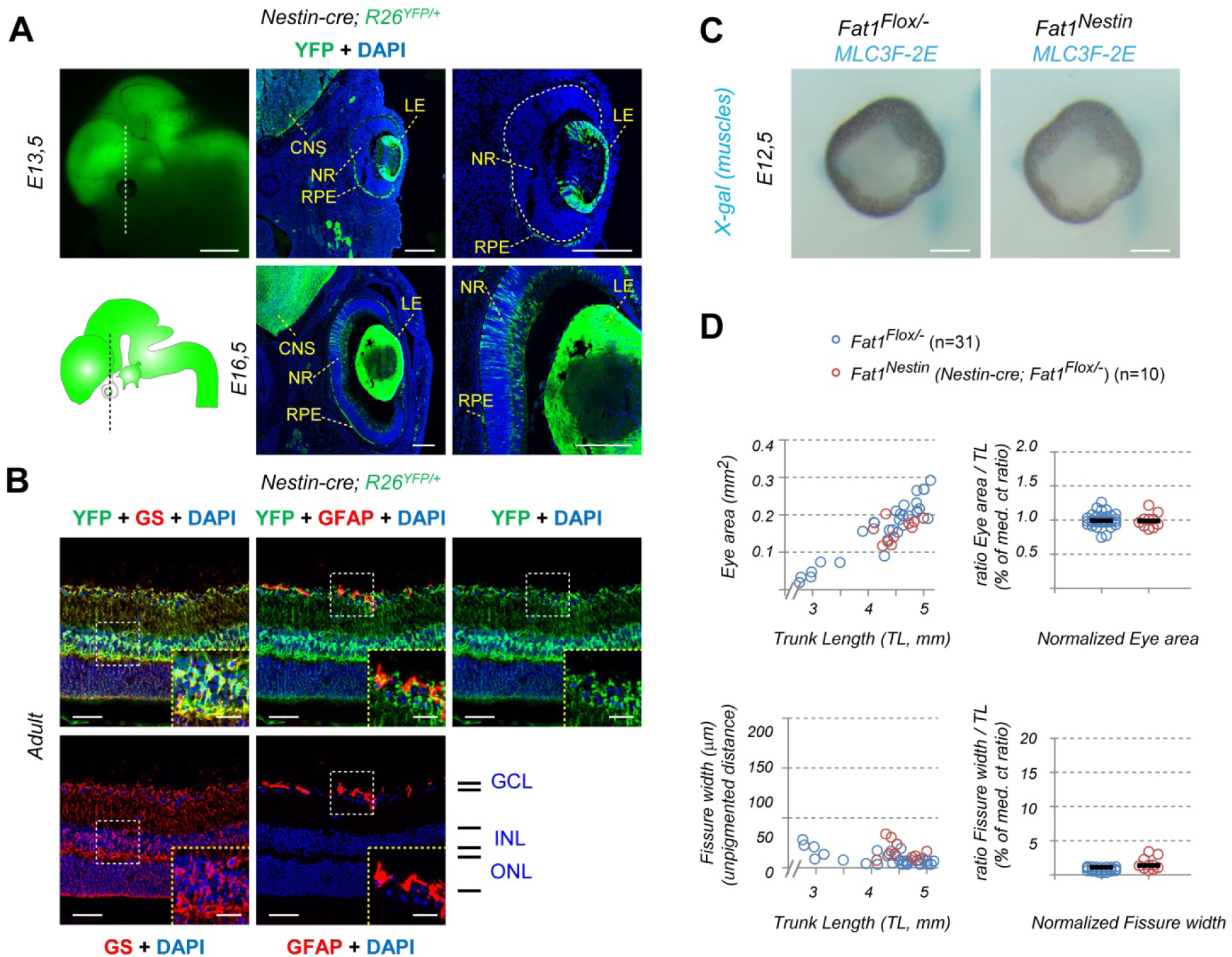
*Nestin-cre*-driven recombination in the neural retina is delayed until after E13.5, thus bypassing stages of optic cup morphogenesis. (A-B) Activity of the *Nestin-cre* line in the developing eye was evaluated at E13.5 (A, upper panels), E16.5 (A, lower panels), or in adult mice (B), by following activation of YFP expression in a *Nestin-cre; R26*^*YFP/*+^ embryo, either by observing direct fluorescence in a whole embryo (A, top left), or by anti-YFP staining on retina sections, combined with DAPI (blue). (A) At E13.5, in contrast to the widespread YFP expression in the nearby thalamus, YFP expression in the eye is restricted to the lens placode, and a small subset of cells in the ventral-most retinal pigmented epithelium, whereas the neural retina is completely devoid of CRE activity. At E16.5, YFP is detected in some cells in the neural retina, among which progenitors and neurons, mostly concentrated in the dorsal-proximal segment, whereas it is detected in virtually all cells in the lens except the center (also devoid of nuclei). (B) In adult *Nestin-cre; R26*^*YFP/*+^ retina sections, YFP immunostaining (green) is combined with antibodies against Glutamine Synthetase (GS, red, left panels) or GFAP (red, middle panels). Inserts show higher magnifications of the regions highlighted by dotted lines, showing cell bodies of Mueller Glia (MG) in the Inner Nuclear Layer (INL, Left panels), or astrocytes in the innermost cellular layer. This analysis confirms that the *Nestin-cre*-derived lineage includes all MG, and most astrocytes. (C) The morphology of the eye was analyzed at E12.5 (top) and E13.5 (bottom) to compare *Nestin-cre; Fat1*^*Flox/-*^ (referred to as *Fat1*^*Nestin*^) with control *Fat1*^*Flox/-*^ embryos (also featuring muscles in blue owing to the *MLC3F-2E* transgene). (D) Quantifications of the width of the ventral optic fissure indicate that the limited and delayed activity of *Nestin-cre* in the neural tissue has fully preserved the functions exerted by *Fat1* to control eye morphogenesis. As a result, *Fat1*^*Nestin*^ embryos exhibit a normally shaped eye, allowing studying the consequences of neural-specific *Fat1* ablation at later stages of retina development. CNS: central nervous system; LE: lens epithelium; NR: neural retina; RPE: retinal pigmented epithelium; GCL: ganglion cell layer; INL: inner nuclear layer; ONL: outer nuclear layer. Scale bars: (A), top left: 1mm; (A) middle and right panels: 200 μm; (B) low mag: 50 μm; inserts 20 μm; (C) 200 μm;

This late CRE activity in the retina thus represents an excellent way of preserving most of *Fat1* expression in the optic cup until completion of optic fissure closure, making it unlikely for *Nestin-cre*-driven *Fat1* deletion to interfere with optic cup morphogenesis, irrespective of the underlying tissue-specificity of *Fat1* activity (and of its proposed requirement in the RPE (Lahrouchi et al., 2019)). This approach indeed allowed preserving *Fat1* function during eye morphogenesis, resulting in normally shaped eyes in which the process of optic fissure closure had been completed without abnormality (Fig. 2C,D). No significant difference in eye morphology was observed between *Nestin-cre; Fat1*^*Flox/-*^ and control *Fat1*^*Flox/-*^ embryos at E12.5 and E13.5, indicating that the *Fat1*-dependent process of optic fissure closure is fully preserved in this genetic setting (Fig. 2C, D). In conclusion, the *Nestin-cre; Fat1*^*Flox/-*^ genetic context (later referred to as *Fat1*^*Nestin*^; see comment in methods) appears suitable to study the role of postnatal and adult *Fat1* expression in the neural retina without interfering with the early process of eye morphogenesis. This tissue-specific approach also allows bypassing non-neural phenotypes with broad systemic consequences such as those occurring in kidney and muscles (Caruso et al., 2013; Ciani et al., 2003; Gee et al., 2016; Helmbacher, 2018). Moreover, it was previously used to demonstrate that depletion of VEGF-A in the *Nestin-cre*-derived lineage resulted in retinal vascular abnormalities caused by dosage-dependent lowering of VEGF-A paracrine signaling (Haigh et al., 2003).

### Fat1 expression in the postnatal retina

We next explored *Fat1* expression in the retina during the phase of postnatal vascular morphogenesis, addressing the relevance of using *Nestin-cre* for *Fat1* ablation. We took advantage of a *Fat1*^*LacZ*^ knock-in allele behaving as a faithful readout of the endogenous *Fat1* expression domain (Caruso et al., 2013; Helmbacher, 2018), and assessed the distribution of *Fat1*^*LacZ*^ expression on either whole mount retinas (Fig. 3A,B, Fig. S1), or retina sections (Fig. 3C, Fig. S1A,B), during the phase of horizontal vascular growth. In flat mounted retinas from P3 *Fat1*^*LacZ/*+^ mice (Fig. 3A,B), endothelial cells, detected with the IB4 lectin, have progressed through one third of the retinal radius (Paredes et al., 2018; Stefater et al., 2011), whereas PAX2-expressing (PAX2^+^) astrocyte progenitor cells (APCs), have nearly reached the retinal periphery (∼80% median progression) (Tao and Zhang, 2014; Tao and Zhang, 2016). β-Galactosidase expression in *Fat1*^*LacZ/*+^ mice was detected throughout the retina, but did not label growing vessels (IB4+), suggesting that at this stage *Fat1* expression occurs in the underlying neural retina, rather than in migrating endothelial cells (Fig. 3B).

**Fig. 3.**
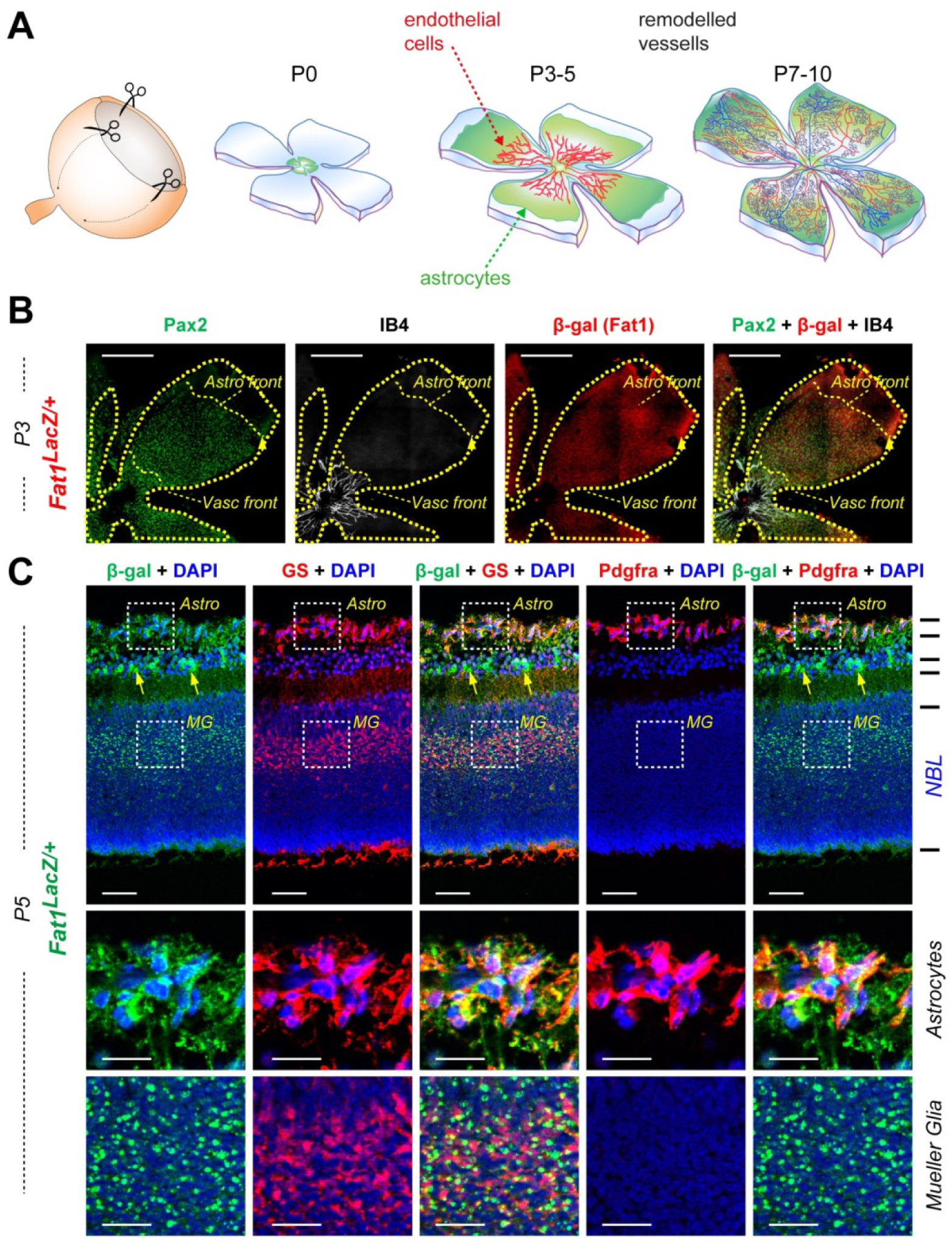
*Fat1* expression in the postnatal retina. (A) Schematic diagrams representing the method for opening retinas for flat-mounting, and the progression of astrocytes and endothelial cells as visualized on flat-mounted retinas. (B, C) *Fat1* expression was followed using the *Fat1*^*LacZ*^ reporter allele in retinas from P3 and P5 mice, and detected in whole mount retinas (B), or on cryosections (C), by immunohistochemistry with anti-β-galactosidase antibodies. (B) Flat-mounted P3 retinas of *Fat1*^*LacZ*^ mice shows a widespread distribution of β-galactosidase expression, not restricted to the regions covered by endothelial cells (identified with IB4 staining, white, the maximum extent being marked as vasc. front), or astrocyte progenitors (identified with PAX2 antibodies, green, the maximum distance being marked as Astro front). (C) Analysis by immunohistochemistry on cross-sections of *Fat1*^*LacZ*^ retinas at P5, with antibodies against β-galactosidase (green), Glutamine Synthetase (GS, red), PDGFRα (red). Cell nuclei were labeled with DAPI. The middle and bottom row panels show high magnification of the regions highlighted with white-dotted areas, focusing on the astrocyte layer (middle row), and an area in the neuroblast layer (NBL), where cell bodies of Müller glia (MG) are found. Scale bars: (B) 500 μm; (C) top row: 50 μm; higher magnifications: 20 μm.

To further characterize in which cell types *Fat1* is present, we immunostained sections of P5 *Fat1*^*LacZ/*+^ retinas, combining β-Galactosidase antibodies with markers of various cell types (Fig. 3C, Fig. S1). As suggested above, we detected a robust β-Galactosidase signal in the so-called neuroblast layer (NBL), in which neurogenesis is still actively occurring (the layering organization being incomplete at that stage). In the NBL, the β-Galactosidase signal closely overlapped with GS signal, mainly marking Müller Glia (Fig. 3C, Fig. S1A,B). In addition, we also detected *Fat1*^*LacZ*^ expression in the innermost retinal layer (RGC layer), in cells exhibiting high levels of PDGFRα and GS (Fig. 3C) or GFAP (Fig. S1B), corresponding to Astrocytes and their progenitors. *Fat1*^*LacZ*^ expression was also apparent in subsets of GS/PDGFRα/GFAP-negative cells located just underneath the astrocyte layer (Fig. 3C, Fig. S1B), likely corresponding to subsets of RGC neurons. In contrast, low *Fat1* levels were detected in endothelial cells (CD31+) and microglia (IBA1+) (Fig. S1A).

*Fat1* is known to be induced in vascular smooth muscle cells (VSMCs) in response to vascular injury to modulate the OXPHOS pathway (Cao et al., 2016), and to cooperate with the transcriptional co-repressor Atrophin to orient VSMC migration (Bruder-Nascimento et al., 2014; Hou et al., 2006; Hou and Sibinga, 2009). *Fat1* was also identified as part of a gene expression signature characterizing arterial endothelial cells (Aranguren et al., 2013). Nevertheless, activation of *Fat1*^*LacZ*^ expression in cells decorating large vessels (putative VSMCs), or in a pattern highlighting arteries at a higher level than veins, was only detected at the end of the period of horizontal growth (P7), and in P21 retinas (Fig. S1C). Thus, *Fat1* expression in VSMCs and arterial ECs arises too late to influence horizontal vascular growth. Consequently, during the period of postnatal angiogenesis, astrocytes represent the likeliest site where *Fat1* expression might be relevant for EC migration.

### Neural Fat1 ablation perturbs astrocyte and endothelial cell horizontal migration

To investigate whether neural *Fat1* activity was required for postnatal angiogenesis, we first analyzed the progression of ECs and astrocytes in retinas from control and *Fat1*^*Nestin*^ pups (Fig. 4). In this situation, non-mutant ECs progress in a *Fat1*-deleted neural territory, with a timing preserving eye morphology (Fig. 2). We first evaluated the extent of vascular progression in P3 and P7 retinas immunostained with CD31 or GS-IB4. Vascular progression was monitored by assessing the distance between the retinal center and the distal vascular front (EC front), relative to the retinal radius (i.e. distance to migrate). In control P3 *Fat1*^*Flox/-*^ mice, the endothelial front had progressed to a median of ∼50% of the retinal radius, whereas *Fat1*^*Nestin*^ mice exhibited a marked delay in vascular growth, with a median progression of 38% of the retinal radius (representing 80% of the progression in controls) (Fig. 4A,C, left plot). At P7, whereas the EC front had reached ∼80% of the vascular radius in control mice, *Fat1*^*Nestin*^ retinas only exhibited a modest but significant shortening of the vascular coverage, with a median progression of 73% (∼95% of the control progression) (Fig. 4B,C, right plot). A delay was also observed in constitutive *Fat1*^*-/-*^ mutant retinas, even when restricting this analysis to quadrants unaffected by the morphological abnormalities associated with coloboma (Fig. S2A,C). Together, our data show that neural-specific *Fat1* deletion transiently delays vascular progression suggesting that neural *Fat1* activity influences the speed/rate of EC migration.

**Fig. 4.**
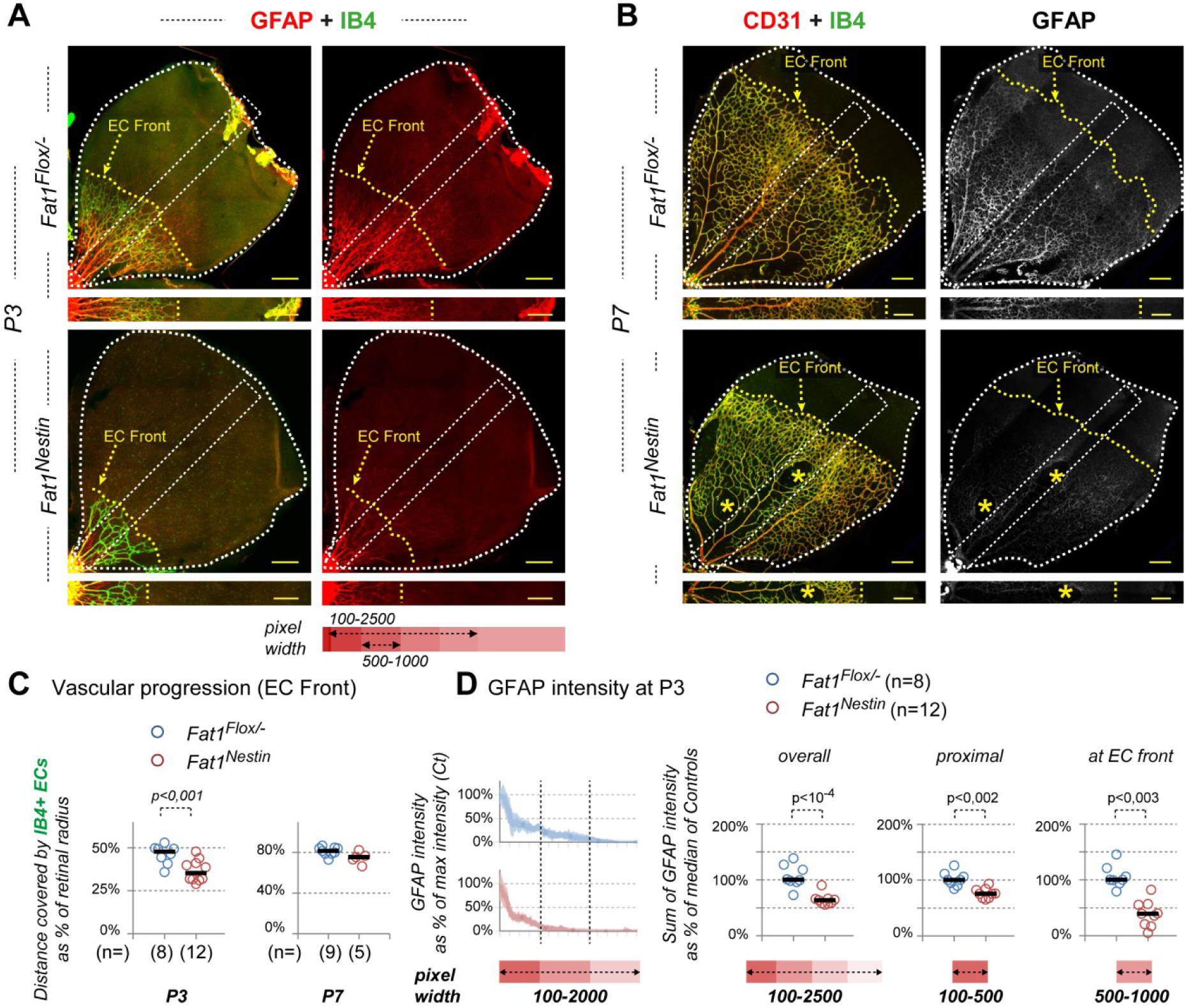
*Fat1* ablation in the neural lineage delays endothelial cell horizontal progression through the postnatal retina. Flat mounted retinas from *Fat1*^*Flox/-*^ control and *Fat1*^*Nestin*^ mutant pups at P3 (A), and P7 (B), after immunohistochemistry with the GS-IB4 lectin (green), an antibody against GFAP (red, A; white, B), and an antibody against CD31 (red, B). Upper pictures feature an entire quadrant of retina. Lower pictures in (A) and (B) correspond to the dotted boxes (extending from the center to periphery of the retinas) in which the quantifications shown in (C and D) were carried out. (C) The distance covered by IB4-positive endothelial cells was measured along the center-to-periphery axis, and expressed as percentage of the retinal radius, at P3 (left plot), and P7 (right plot). (D) The intensity of GFAP staining was quantified along the radius of each retina (in dotted areas), and the average intensity distribution plotted for each genotype (D, left plots). The sum of signal intensity was calculated for a window spanning the distances schematically represented below the images in (A), the width of which is indicated (in pixel coordinates) below each plot (D), covering either the full width of endothelial layer (left plot, 100-2500), a small window proximal to the front of migration (middle plot, 100-500, even in mutant retinas), or spanning the migration front (right plot, 500-1000). Scale bars: (A, B) 200 μm.

Given the pattern of CRE activity, the delayed horizontal progression of ECs observed in *Fat1*^*Nestin*^ retinas unambiguously represent a non-cell-autonomous consequence of neural-*Fat1* ablation. We therefore asked whether this phenotype was associated with alterations in the distribution of cell types included in the *Nestin-cre* lineage and involved in angiogenesis. Among those, astrocytes and astrocytes progenitors play an active role driving endothelial cell migration (Selvam et al., 2018; West et al., 2005), and expressed robust *Fat1* expression (Fig. 3B). Astrocyte distribution and maturation can be followed with the astrocyte marker GFAP. GFAP levels are low in immature astrocytes (in the EC-free distal portion), and high in mature astrocytes (in the EC-covered region), reflecting their switch from immature/proliferative to mature phenotype (Rattner et al., 2019; Tao and Zhang, 2014; West et al., 2005) (Fig. 4). We therefore analyzed GFAP distribution to determine whether astrocyte distribution/maturation was dependent on *Fat1* activity. The sharp transition in the intensity of GFAP signal coincides with the front of endothelial cell migration (Fig. 4A,B; dotted line). As expected, the position of the transition between GFAP_high_ and GFAP_low_ levels matched the degree of vascular progression, in control and mutant contexts, reproducing the delay observed in *Fat1*^*Nestin*^ retinas, robust at P3, and milder at P7 (Fig. 4A-C, Fig. S2A). Nevertheless, even in the region covered by endothelial cells, GFAP intensity appeared lower in mutant retinas. We therefore used image processing to assess the effect on GFAP signal intensity, measured GFAP intensity along a window spanning from the retina center to the periphery, and calculated the sum of intensity. This analysis confirmed the global lowering of GFAP intensity throughout the retinal radius (encompassing both vascularized and non-vascularized areas), not only at P3 (Fig. 4D, left plot), but also at P7 (Fig. S2D). A large part of this reduction may be accounted for by the delay in EC migration, since astrocyte maturation (i.e the sharp increase in GFAP level at the EC transition) is driven by the contact with endothelial cells and their sensing of the oxygen supply associated with incoming blood (Duan et al., 2017; Duan et al., 2014; West et al., 2005). We therefore restricted this analysis to a small window, proximal to the vascular front (outlined by a yellow dotted line in Fig. 4A,B), where all astrocytes, even in mutant contexts, are in contact with ECs. This also uncovered a strong lowering in GFAP levels in *Fat1*^*Nestin*^ retinas compared to controls (Fig. 4D, middle plot, window 100-500). Thus, there was a persistent effect on astrocyte maturation even at a distance from the vascular front (Fig. 4D, right plot), indicating that *Fat1* activity is required for astrocyte maturation even in the central-most astrocytes, after they underwent the EC-induced transition.

### Neural Fat1 ablation destabilizes the astrocyte layer

Although vascular progression is only delayed, with a relatively modest residual effect at P7, we consistently observed patches devoid of vascular cells in the rear of the EC migration front of P7 *Fat1*^*Nestin*^ (asterisks in Fig. 4B, 5A), or as large indentations in the EC front in *Fat1*^*-/-*^ retinas Fig. 5A, S2A). Such patches were not observed in control retinas and were characterized by a lack of both endothelial cells and astrocytes, as indicated by the match in shape between CD31 and GFAP_high_ distribution (Fig. 4B, 5A, S2A, asterisks). When observed in the vascularized area, these patches appeared rounded, irrespective of their size and position, as if resulting from a rupture in the EC/astrocyte layer, followed by tissue-retraction. The underlying retinal structure was preserved, suggesting that these patches specifically resulted from interruptions of the layer containing astrocytes and endothelial cells. Thus, even if the effect on migration dynamics of endothelial cells and astrocytes is transient, neural-*Fat1* inactivation also leads to local dissociation of the EC/Astrocyte layer from the underlying neural retina, potentially indicating a compromised adhesion between the astrocytes and underlying Müller glia.

**Fig. 5.**
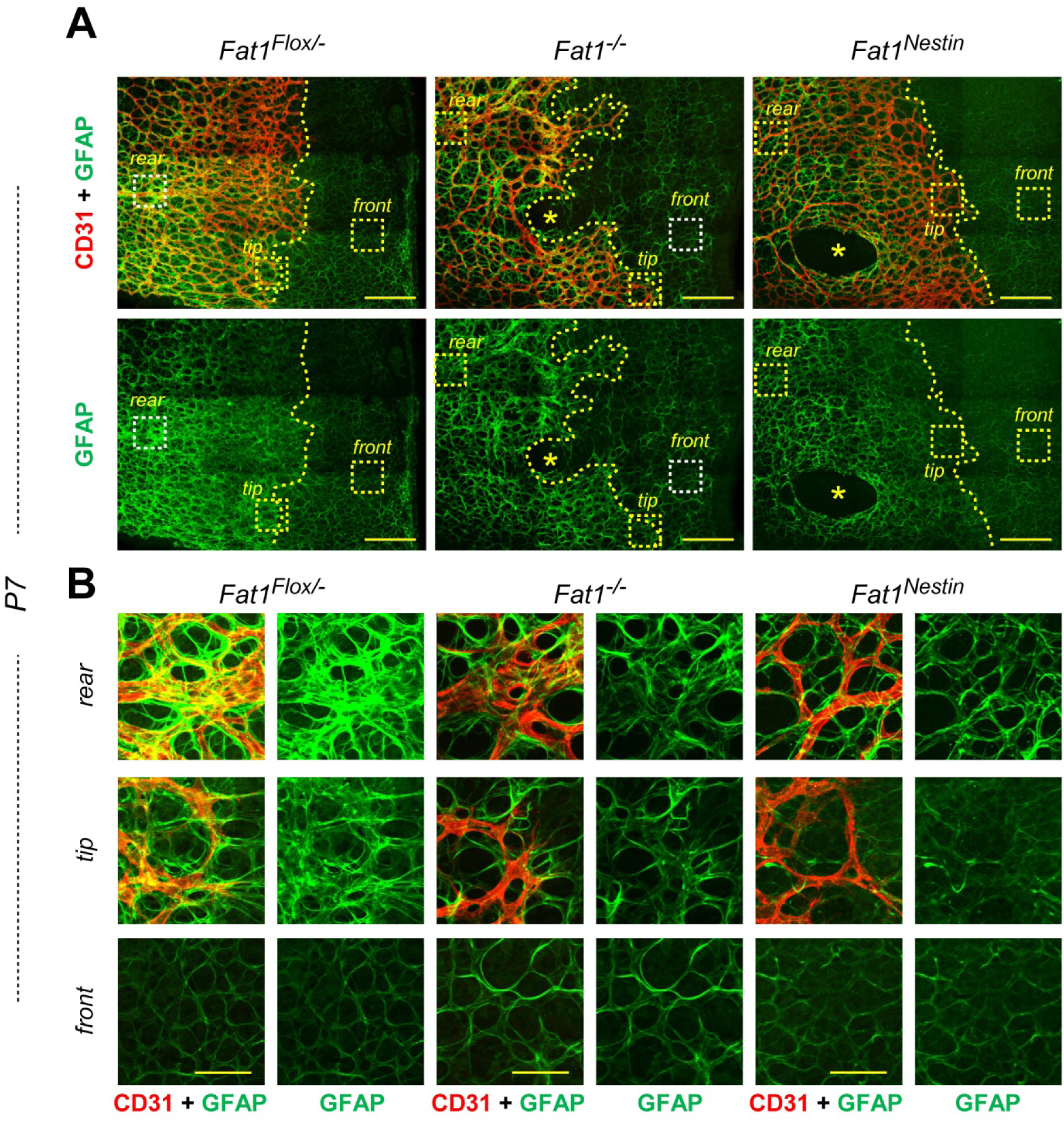
*Fat1* ablation in the neural lineage alters astrocyte maturation. Flat-mounted retinas from *Fat1*^*Flox/-*^ control, *Fat1*^*-/-*^ and *Fat1*^*Nestin*^ mutant pups at P7 after immunostaining with antibodies against CD31 (red) and against GFAP (green). Upper pictures (A) represent an area of the retina spanning from near the center (to the left), where the astrocyte layer is covered by endothelial cells, to the distal most part of the retina (to the right), where GFAP levels are low in absence of vascular coverage. Lower pictures in (B) correspond to the dotted boxes highlighted in (A), visualizing for each genotype, a region 1) in the vascularized area (rear); 2) at the level of tip endothelial cells (tip); and 3) in the “avascular” distal region (front). Scale bars: (A) 200μm, (B) 50μm.

We next analyzed astrocyte coverage at higher resolution. The sharp boundary in GFAP level seen at low magnification (Fig. 4) corresponds, at higher resolution, to an increase in the density of the GFAP^+^ fibrillary meshwork in areas where astrocytes are in contact with endothelial cells, compared to areas devoid of endothelial cells (also called hypoxic or pre-vascular areas) (Fig. 5). This analysis confirmed that the lowering in GFAP levels in *Fat1*^*-/-*^ and *Fat1*^*Nestin*^ retinas reflected a lower fibril density, and uncovered irregularities in the fibrillary meshwork (Fig. 5A, 5B). Such changes in GFAP coverage were observed 1) in the vascularized areas (in mature astrocytes, in proximal areas in the rear of the EC front; First row in Fig. 5B), 2) at the level of the EC migration front (tip cells, where astrocytes undergo their transition; Second row in Fig. 5B), and 3) in distal regions ahead of the vascular front (in pre-vascular APCs (third row in Fig. 5B)). In these pre-vascular areas, the meshwork of GFAP^+^ fibrils delimits non-stained spaces, which appeared larger or more irregular in size in mutant retinas (*Fat1*^*-/-*^ and *Fat1*^*Nestin*^; Fig. 5B, distal). The presence of such morphological alterations in pre-vascular astrocyte-progenitors rules out the possibility that defects in maturation would exclusively be secondary to delayed EC migration, thus reinforcing the possibility that *Fat1* activity directly influences astrocyte phenotype, even prior to their Oxygen/EC-induced transition.

### Neural Fat1 ablation delays astrocyte maturation

To complete our characterization of defective astrocyte maturation, we next examined expression of PDGFRα and PAX2 in P3 retinas (Fig. 6). PDGFA is a key contributor that feeds astrocyte progenitor cell migration: Secreted by retinal ganglion cells, PDGFA acts on APCs and immature astrocytes expressing its receptor PDGFRα, to promote their migration (Fruttiger et al., 1996; Gerhardt et al., 2003; Tao and Zhang, 2016). Similar to GFAP staining, PDGFRα levels are correlated with the progression of astrocyte maturation (Fig. 6A-C). However, the increase in PDGFRα expression associated with maturation is not as abrupt as that of GFAP, but occurs though a steady slope between the APC migration front and the vascular front, with lowest levels in distal APCs, and highest levels observed in mature astrocytes in the vascularized/proximal retina (Fig. 6A-C). To evaluate changes in PDGFRα levels resulting from neural *Fat1* ablation, we analyzed staining intensity across the retinal radius using the same method as used earlier (Fig. 6C), and plotted the sum of intensity corresponding to either the full span or windows corresponding to pre or post-vascular areas (Fig. 6D). This analysis confirmed not only the overall lowering of PDGFRα levels in *Fat1*^*Nestin*^ retinas compared to controls (Fig. 6D, left plot), again consistent with the delayed EC migration dynamics, but also a robust lowering at the level of the transition zone (Fig. 6D, middle segment) and in the pre-vascular zone (Fig. 6D, distal segment), consistent with a flattening of the slope of PDGFRα levels in non-vascularized astrocytes. In contrast, no significant change was observed between *Fat1*^*Nestin*^ and control retinas in the proximal-most region, where PDGFRα levels plateau at their highest level in mature astrocytes (Fig. 6D, proximal segment).

**Fig. 6.**
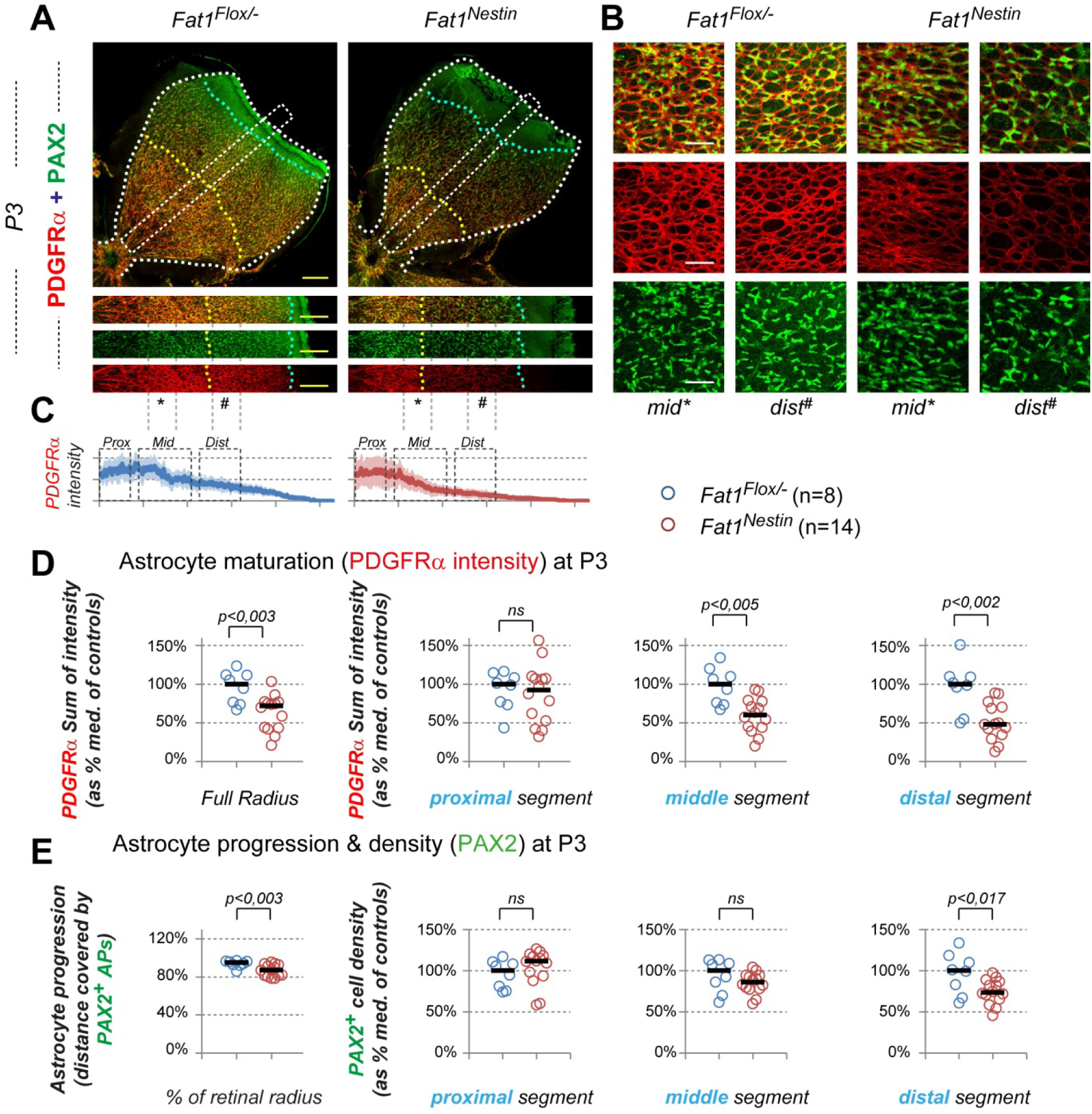
Fat1 ablation in the neural lineage interferes with migration and maturation of astrocyte progenitors. Flat-mounted retinas from *Fat1*^*Flox/-*^ and *Fat1*^*Nestin*^ mice at P3 stained with PDGFRα (red), and PAX2 (green) antibodies. (A) Upper pictures feature an entire quadrant of retina, while the lower pictures correspond to the dotted boxes (extending from the center to periphery of the retinas) in which the quantifications shown in (C-E) were carried out. (B) High magnification images corresponding to the zones within the dotted boxes, highlighted by the signs * and ^#^, corresponding to the areas defined as middle and distal in the quantifications. Because of the mild shortening of the distance covered by astrocytes, we purposely positioned the “distal^#^” area posterior to the astrocyte front, even in mutant retinas. (C) PDGFRα signal intensity was measured along the center-to-periphery axis, and plotted along the proximo-distal axis. (D-E) Quantitative analyses of the Sum of PDGFRα intensities (D), or of the extent and density of PAX2^+^ astrocytes (E). (D) For each retina, the sum of PDGFRα intensity (expressed as percentage of the median value of controls) was calculated for either the full radius (left plot), or for segments outlined in (C), defined as proximal, middle and distal. (E) Plots represent (as percentage of the median value of controls), the distance reached by PAX2^+^ astrocytes (left plot), or the density (number of PAX2^+^ cells per area) in each of the defined segments (proximal, middle, and distal). Scale bars: (A): 200μm; (B) 50μm.

By highlighting nuclei of migrating APCs and immature astrocytes (Duan et al., 2014; Tao and Zhang, 2014; Tao and Zhang, 2016), PAX2 staining allows evaluation of changes in cell number and density, and in the distance travelled through the retina. At P3 in control retinas, the overall distance covered by astrocytes (between the retina center and the front of astrocyte migration) represented nearly the full radius (∼90%, Fig. 6A,E). Although modest, there was a significant shortening of the distance covered by PAX2^+^ cells in mutant P3 retinas of *Fat1*^*Nestin*^ mice (Fig. 6E, left plot). Nevertheless, the reduction in PDGFRα levels described above (Fig. 6D) largely occurred in areas where we did not detect significant changes in the density of PAX2^+^ cells (Fig. 6E, density plot in *middle* segment). This indicates that the lowering in PDGFRα levels is not the mere consequence of a reduced astrocyte density, but a true effect on protein levels, hence on astrocyte maturation. Nevertheless, the mild shortening of the astrocyte extent was associated with a significant lowering in the density of PAX2^+^ cells in pre-vascular (distal) regions (Fig. 6E, distal segment). Although we haven’t ruled out a possible change in AP proliferation, these observations are indicative of an abnormal migration process. The latter effect is consistent, given the known role of PDGFRα signaling on astrocyte migration, with a consequence of lowered PDGFRα level in the distal-most migrating APCs.

Given the known role of FAT Cadherins in modulating polarized migration events, we next asked if this effect of *Fat1* deficiency on APC migration and density in distal segments were associated or preceded by alterations in cell polarity. Owing to the PAX2 nuclear labelling, we noticed that in control retinas, the nuclei of astrocyte progenitors exhibited a strong orientation preference representing the main radial direction (Fig. 7). This was particularly manifest near the migration front, where PDGFRα levels are the lowest, whereas in the rear, PAX2^+^ nuclei were aligned along the cellular connections highlighted by PDGFRα staining (Fig. 6B). We therefore measured for individual PAX2^+^ APCs, the angle between the main direction of migration (represented by the radius in a band of limited width), and the vector formed by the longest nuclear length. When performed on all APCs present in the distal-most segment at the migration front, this revealed the percentage of nuclei in each defined angle-range per mouse (Fig. 7C). Whereas the highest represented angle-range in control mice was the outward direction (range 80-100°), neural-*Fat1* ablation led to a marked change in angle distribution, with a lower proportion of nuclei matching the outward direction, and increased percentages found in the (pooled) 0-60 and 120 to 180° ranges (Fig. 7C).

**Fig. 7.**
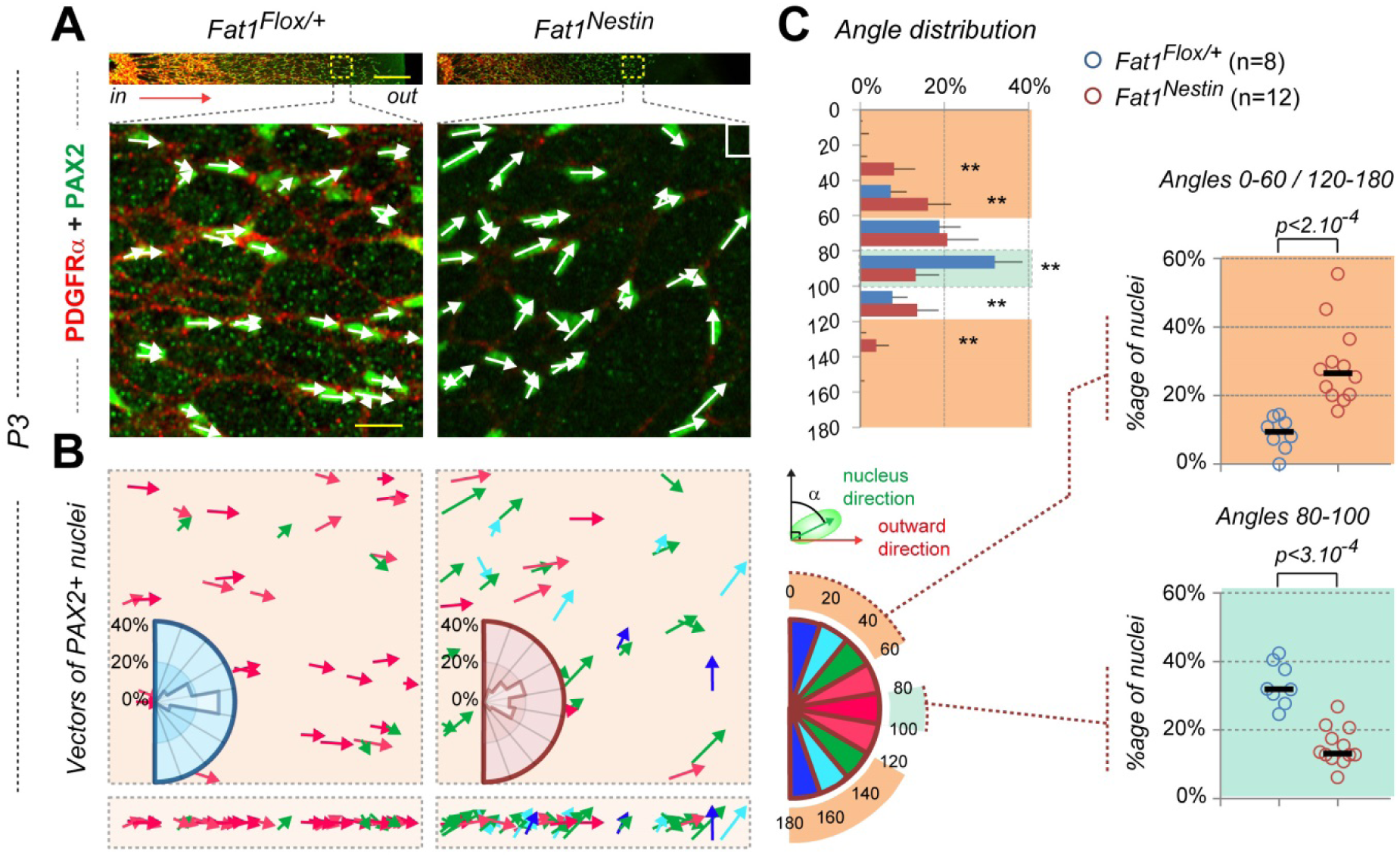
Fat1 controls the polarity of astrocyte progenitor migration. Imaging of astrocyte progenitor polarity in flat-mounted retinas from *Fat1*^*Flox/-*^ and *Fat1*^*Nestin*^ mice at P3 stained with PDGFRα (red), and PAX2 (green) antibodies, as in Fig. 6. (A) Upper pictures represent a region extending from the center to periphery of the retinas, corresponding to the white dotted boxes in Fig. 6A. The lower pictures are higher magnification images of astrocyte progenitors near the migration front corresponding to the squared dotted boxes in the bands above, in which the quantifications shown in (B, C) were carried out. A white arrow was placed on top of each PAX2^+^ astrocyte progenitor to highlight the directional vector of its longest dimension (nucleus length). (B) For each genotype, the vectors corresponding to the examples above were color coded as defined on the color-rose plot on the right, and a polar plot shows the angle distribution of the PAX2^+^ vectors. The main migration direction (from the center to the periphery) in the imaged area was set to 90°. Polar plots are representative of *Fat1*^*Flox/-*^ control (n=8), and *Fat1*^*Nestin*^ mutant (n=12) mice, with ∼80-100 cells measured per mouse at the migration front (where PDGFRα expression level is the lowest). (C) Left plot: Alternative representation plotting the overall angle distribution of PAX2^+^ vectors, showing the proportion of cells in each angle range (error bars = standard deviation; **: statistical significance, p<0.001). Right: scatter plots showing the proportion of PAX2^+^ astrocytes exhibiting angles in the 80-100°, and 0-60°/120-180° angle ranges, respectively (the first category representing the preferential direction in controls). Scale bars: (A) top: 200μm; bottom: 20μm.

Together, our results indicate that *Fat1* deletion interferes with migration of astrocyte progenitors and with their maturation into immature astrocytes. One of the earliest manifestations is a loss of the preferential APC nuclei orientation (normally oriented outward), immediately followed by a failure to upregulate PDGFRα expression. Both changes impact the rate of astrocyte migration, and consequently the rate of endothelial cell progression in the postnatal retina. Although these effects are transient, *Fat1* deletion also compromises adhesion between astrocytes and the underlying Müller glia.

### Neural-specific Fat1 ablation leads to abnormal architecture of adult retinal vasculature

To evaluate the consequences of postnatal phenotypes on adult retinas, we next explored the anatomy of the vascular network in control and *Fat1*^*Nestin*^ mutant retinas, by analyzing the layered structure of the retinal vasculature after immune-labelling of whole mount retinas with endothelial markers (CD31, shown in Fig. 8, but also GS-IB4 and Claudin5, shown in Fig. 9). Retinas were flat mounted with their inner side (vitreal side) on the top, and we imaged either a whole retina quadrant at low resolution, or the distinct layers at higher resolution (Fig. 8A). The adult retinal vascular network is organized in three layers, intercalated with neuronal layers (Fig. 8B-D). The large vessels (arteries and veins) radiate through the plane of the retina, along its inner side, from the center to the periphery, sharing space and direction with axons of retinal ganglion cell (RGC) neurons, forming the so-called nerve fiber layer (NFL) (Ruhrberg and Bautch, 2013). This innermost NFL is thus constituted of the veins and arteries, and of their first order ramifications. These first order branches further ramify into smaller branches with diameter decreasing with the hierarchical level, and plunge deeper in the retina via vertical branches. At two distinct retinal levels, respectively corresponding to the interface between the inner nuclear layer (INL) and outer nuclear layer (ONL) and between the ONL and the rods and cone layer (Fig. 8C), these vessels make 90° turns, and establish two honeycomb-like horizontal vascular networks (mostly composed of capillaries), respectively called the inner, and outer plexiform layers (IPL and OPL, Fig. 8C,D). Once retinas are flat-mounted, it is possible to acquire images of fluorescent labeling throughout the depth of the retina (without the need for specific clearing method), and to process separately the focal planes corresponding to the three layers. Thus, 3 images of fixed areas are either sequentially acquired or extracted from a stack to visualize the 3 vascular levels for a same position in the retina (Fig. 8B,D; indicated by a yellow-dotted square in Fig. 8A). The three layers are then assigned a different virtual color, and merged as a flat image of to reconstitute the whole vasculature in the area (Fig. 8B, bottom row).

**Fig. 8.**
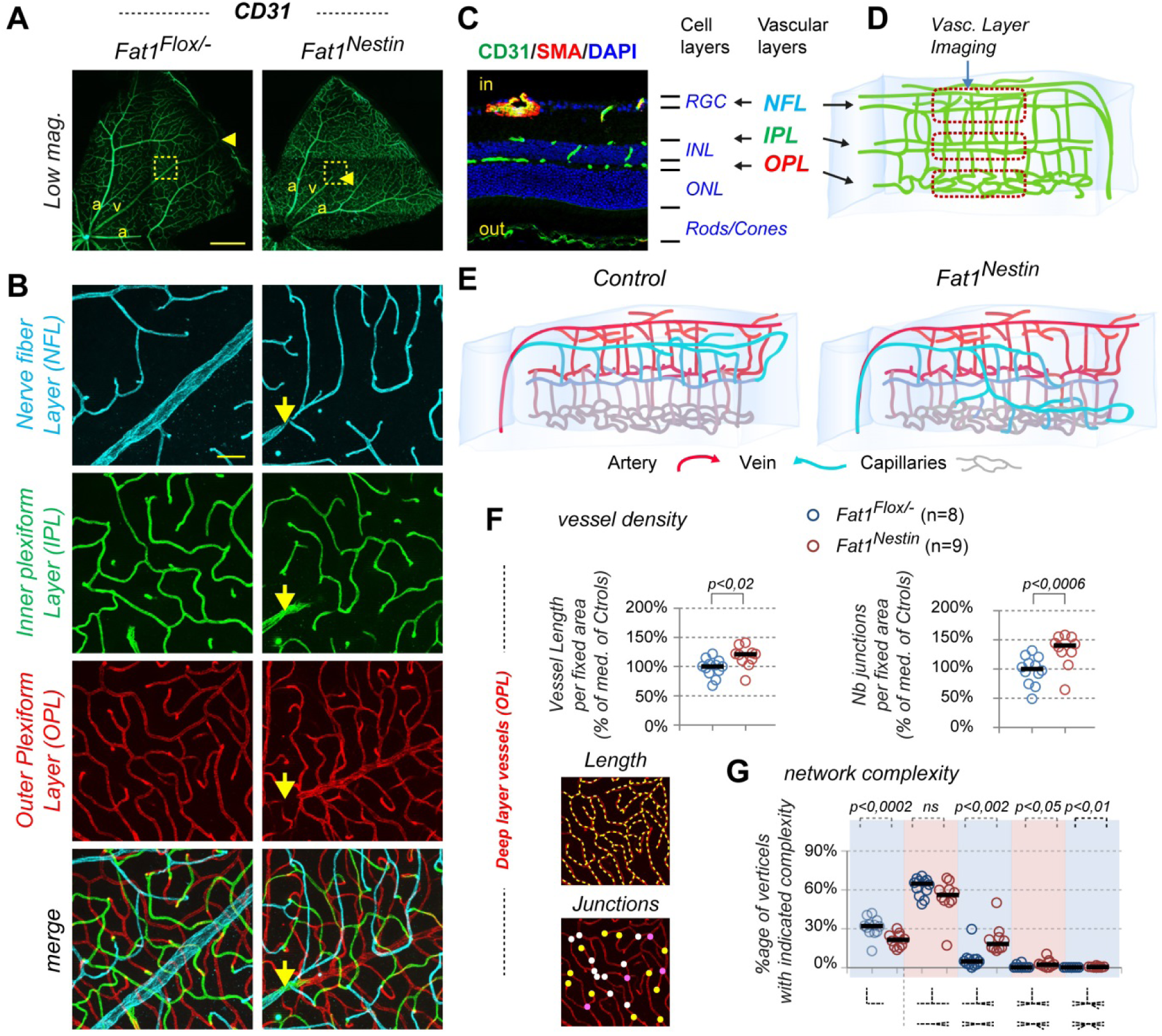
Adult mice with neural Fat1 ablation model vascular pathology. (A, B) Images of flat mounted retinas from *Fat1*^*Flox/-*^ control and *Fat1*^*Nestin*^ mutant adult mice, immunostained with CD31 antibodies, visualizing either one retina quadrant (A), or high magnification images of the 3 vascular layers in the squared dotted regions (B), with each of the 3 images corresponding to Maximal Intensity projections (MIP) of optical stacks as schematized in (C), corresponding respectively to the Nerve fiber layer (NFL), Inner plexiform layer (IPL), and outer plexiform layer (OPL. MIPs of each layer were assigned different colors, and merged (bottom panels). (C) Section of an adult retina immunostained with CD31 (green), αSMA (Acta2, red), and DAPI, to illustrate the position of vascular and nuclear layers. (D) Schematic representation of the vascular organization in flat-mounted adult retinas (from control mice) showing (center to the right) the levels within the retinal depth corresponding to the layer images shown in (B). (E) Schematic representation of the distribution of vessel types (with arteries and arterioles in red, veins and venules in blue, and capillaries in grey) and their distribution in vascular layers in control (left) and in *Fat1*^*Nestin*^ (right) retinas. (F, G) Quantitative morphometric analysis of changes in the vascular architecture of deep retinal layer vessels caused by neural *Fat1* ablation: (F) Plots representing the change in total vessel length per area (left plot), and number of junctions (right plot) measured in defined area, normalized to the median value of control mice (set to 100%), each dot representing one mouse. (G) Vascular network complexity is evaluated by quantifying the number of junction between branches (vertical/horizontal) associated with each intersection in the deep vascular layer. The plot represents (for each genotype, one dot per mouse), the percentage of intersections with each complexity range, as represented schematically below the graph. Scale bars: (A) 500μm; (B): 50μm.

**Fig. 9.**
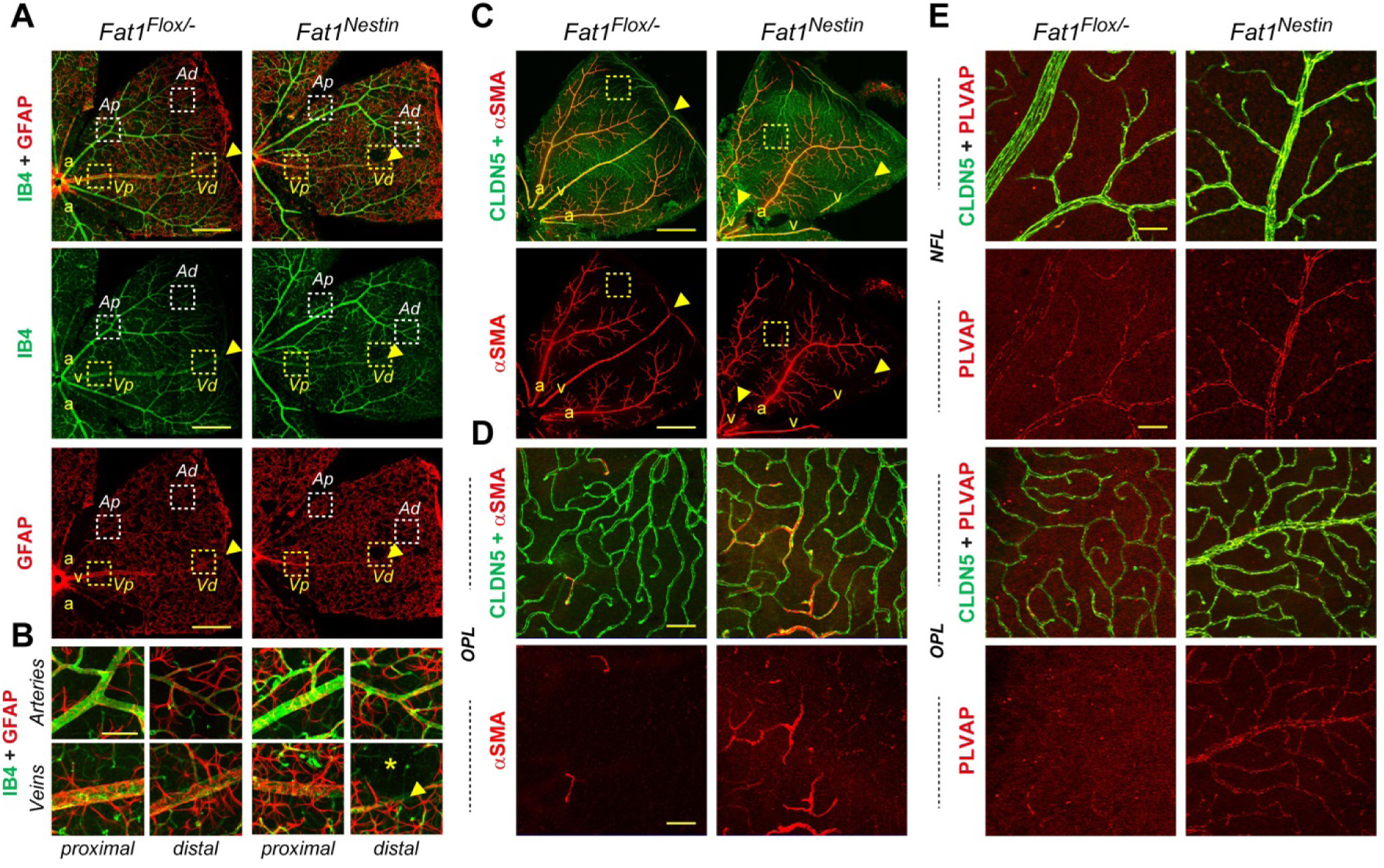
Adult mice with neural Fat1 ablation exhibit disrupted blood brain barrier. (A) Images of flat-mounted retinas immunostained with: (A, B) IB4 (green) and GFAP (red); (C, D) Claudin5 (Cldn5, green) and αSMA (Acta2, red); (E) Cldn5 (green) and PLVAP (red), visualizing either a quadrant (A, C), or regions highlighted in (A, C) with yellow-dotted squares. Scale bars: (A, C) 500μm; (B): 100μm; (D, E): 50μm.

Overall, the layered structure of retinal vasculature and the global vessel hierarchy were relatively preserved in adult *Fat1*^*Nestin*^ retinas. Nevertheless, we noticed on mosaic images of retina quadrants that *Fat1*^*Nestin*^ retinas exhibited a different relative distribution of veins and arteries (schematized in Fig. 8E). In control retinas, veins typically form by convergence of collecting vessels towards the edge of the retina, where they coalesce into larger veins, which in turn fuse (arrowhead, Fig. 8A,B) to form one large vein directly returning to the center to exit the retina through the optic stalk. In contrast, in *Fat1*^*Nestin*^ retinas, we frequently observed that only the proximal portions of large veins could be distinguished in the NFL (starting from a discrete point found in random position along the retinal radius, arrowhead in mutant retina, Fig. 8A,B). When imaging the 3 layers around such a point, we could observe a transition between a collecting vessel running in the deep OPL layer, suddenly turning upward and emerging in the upper layer (Fig. 8A,B,E).

To measure the impact of this change, we used a quantitative morphometric approach. We evaluated vessel density in the three layers, by measuring in fixed areas for each of them, the cumulated vascular length (see methods), and the number of intersections (representing either intersections between vertical and horizontal branches, or junctions of several horizontal branches) (Fig. 8F). This method uncovered an increased vascular density (with increased cumulated length and number of intersections) in the deep layer (OPL) of *Fat1*^*Nestin*^ retinas compared to controls (Fig. 8E). Furthermore, the complexity of branching patterns was also significantly increased in the OPL of *Fat1*^*Nestin*^ compared to *Fat1*^*Flox/-*^ control mice (Fig. 8G), with more intersections representing junctions of larger numbers of vascular segment (junctions of 4, 5 or 6 segments), and less junctions associating one vertical and one horizontal segment. Overall, this analysis revealed that *Fat1* ablation in the neural lineage leads to the emergence of large collecting vessels in deep vascular layers, associated with an increased vascular density.

Given the postnatal phenotype in astrocytes described above, we next wondered whether the observed alterations of the adult vascular architecture might be correlated with changes in astrocyte distribution. Starting from the stage when astrocyte migration is completed, the vascular remodeling leading to the appearance of higher order vessels is associated with a redistribution of astrocytes in tight association with large vessels in the NFL (Fig. 4B), with a strong bias towards veins in adult retinas (Fig. 9A). Thus, in adults, veins are distinguishable from arteries by their higher astrocytic coverage, as well as by their lower intensity of IB4-binding (Fig. 9A). Interestingly, we noticed that the point of transition described above, where veins shift from deep layers to the NFL in *Fat1*^*Nestin*^ mutant retinas (yellow arrowheads in Fig. 8A, 9A,B; arrows in Fig. 8B) was marked by a hole in the astrocyte coverage (Fig. 9A, asterisk), reminiscent of the irregularities observed in P7 retinas. We cannot distinguish whether these astrocyte-free zones are the same as those observed in postnatal retinas, which would have persisted till adulthood, or whether they have formed de-novo.

### Neural Fat1 ablation alters vascular integrity

Intrigued by the increase in vascular density and complexity in deep layers of *Fat1*^*Nestin*^ mutants, we next asked whether we might identify other signs that deep vessels, which are normally exclusively capillaries, might have acquired characteristics of higher order vessels. One such characteristic is the presence of vascular smooth muscle cells (VSMCs), identified by expression of Smooth muscle actin α (αSMA, also referred to as Acta2 for actinα2) (Fig. 9B). Mosaic images of flat-mounted retinas show that large vessels are strongly decorated with VSMCs, which are also present on first order arterial ramifications (Fig. 9B). Instead, imaging of deep vascular layers (in yellow-dotted squares), reveals that deep capillaries are almost completely devoid of VSMCs in control retinas. In contrast, in mutant retinas, we detected spots in the OPL where many capillary segments were decorated with VSMCs, even in areas in which they do not show high degree of complexity associated with collecting vessels (as in Fig. 8B, 9C). Thus the increased vascular complexity of deep layers is associated with VSMC recruitment. Given that VSMCs do not derive from the *Nestin-cre* lineage, this phenotype represents an additional non-cell autonomous consequence of *Fat1* depletion.

Finally, investigated the expression of PLVAP, an adhesion molecule characteristic of vessels with endothelial fenestrations, the silencing of which being necessary to ensure blood brain barrier integrity in central nervous system vessels (Shue et al., 2008; Stan et al., 2004). Upregulated PLVAP labelling was observed in 2 out of 5 *Fat1*^*Nestin*^ mutant retinas analyzed, but not in control retinas (Fig. 9C). This increase, however, was not associated with a lowering of the tight junction protein CLDN5 (Fig. 9B, 9C), also considered key to ensure high endothelial barrier function. Thus, the non-cell autonomous consequences of *Fat1* ablation in the neural lineage can lead in some cases to molecular changes typical of altered blood brain barrier integrity. This implies that *Fat1* activity in the neural lineage (including astrocytes and Müller glia), is necessary for the maintenance of vascular integrity and barrier function.

## Discussion

In this study, we investigated how the FAT1 Cadherin modulates retinal vascular development and integrity. As *Fat1* is required for optic cup closure during development, we bypassed this early requirement by disrupting its activity in the late neural linage owing to late-onset of *Nestin-cre* activity in the retinal epithelium, thus preserving eye morphogenesis. In this context, we found that *Fat1* expression in the neural lineage, predominantly occurring in astrocytes and Müller glia, is required for coordinated migration polarity and subsequent maturation of astrocyte progenitors in the postnatal retina. Alteration of both processes in *Fat1*^*Nestin*^ mutants consequently delays retinal angiogenesis, and also results in stochastic detachment of astrocytes from the underlying layers. Some abnormalities persisted at adult stages, as measured through an increase in the density of deep vascular layers and changes in distribution of higher order vessels. Thus, *Fat1* activity in the neural lineage is required for polarized migration and maturation of astrocyte progenitors, and its deletion interferes with the astrocyte/endothelial cell cross-talk, thus delaying retinal angiogenesis.

### Mechanisms of astrocyte maturation and coupling with angiogenesis

Our study has uncovered a key role of FAT1 signaling in astrocyte maturation, the suppression of which affects both the transition from astrocyte progenitors to immature astrocytes, ahead of the vascular migration front, and further maturation of astrocytes in vascularized retina. Whereas the latter aspect is likely a complex consequence of the delayed progression of endothelial cell migration, the first aspect represents a direct consequence in astrocytes of *Fat1* deficiency, manifested by a lowering of PDGFRα protein levels (Fig. 6). PDGFRα expression gradually increases in migrating astrocyte progenitors between the front of migration and the EC front, tuning their sensitivity and responsiveness to PDGF produced by the underlying RGC neurons. One of the functions assigned to PDGF/ PDGFRα signaling in astrocytes is to promote their migration (Fruttiger et al., 1996). Inhibition of PDGFRα signaling in the postnatal retina, by infusion with function blocking antibodies (Fruttiger et al., 1996) or by *Pdgfrα* ablation in the astrocyte lineage (Tao and Zhang, 2016), was shown to abrogate migration of astrocyte progenitors, and to consequently suppress the attractive VEGFA gradient for ECs, thereby blocking their horizontal migration (Fruttiger et al., 1996; Tao and Zhang, 2016). In contrast, gain of PDGFRα signaling (in transgenic mice overexpressing PDGF-A) induced an increase in astrocyte and vessel density, nevertheless also slowing migration (Fruttiger et al., 1996). The flattened slope of PDGFRα increase observed in *Fat1*-deficient astrocytes implies that PDGFRα expression level is regulated by FAT1 signaling. Regulation of PDGFRα by *Fat1* might either occur directly in astrocytes, or may represent a non-cell-autonomous consequence in astrocytes of *Fat1* activity in interacting cell types, such as Müller glia or neurons.

Interestingly, astrocyte progenitors at the migration front exhibit a marked elongation of their nuclei in the direction of migration aligned along the retinal radius, and *Fat1* disruption resulted in a loss of this preferential orientation. Together with the reduced PDGFRα levels discussed above, this led to a delay in astrocyte migration, and to a reduced astrocyte density restricted to the front of astrocyte progression. Thus, retinal astrocytes represent a novel model of planar and polarized migration, the modulation of which is functionally linked with the dynamics of angiogenesis. This preferential direction is instructed, at least in part, by radially oriented RGC axons (O’Sullivan et al., 2017). The polarization is best seen at the migration front, where PDGFRα levels are the lowest, whereas in the rear, nuclei are aligned along PDGFRα^+^ cellular processes extending between cells. The change in polarity caused by *Fat1* deficiency is detected in these distal-most APCs. The co-occurrence of both phenotypes raises the possibility that the control of PDGFRα levels by *Fat1* might be an indirect consequence of the altered migration polarity, rather than a direct transcriptional or translational outcome of FAT1 signaling. Distinguishing between the two possibilities will necessitate an experimental approach allowing uncoupling of the two phenomena, so as to determine whether PDGFRα levels are regulated in leading APCs by tension forces associated with cell polarity, or in follower APCs by the traction forces exerted by leading APCs.

The astrocyte migration/maturation phenotype was associated with a robust delay in the rate of endothelial cell migration across the retina. Astrocyte maturation and endothelial cell migration are indeed tightly coupled. A key contributor in this coupling is the oxygen supplied by blood perfusion of the newly formed vessels, which converts immature astrocytes into mature astrocytes (Duan et al., 2017; Duan et al., 2014; West et al., 2005), in part by promoting GFAP expression, and by repressing VEGF expression, otherwise induced by hypoxia (Duan et al., 2017; Duan et al., 2014; Rattner et al., 2019; West et al., 2005). Aside from oxygen, other EC-derived factors also contributing to astrocyte maturation include ApelinJ (Sakimoto et al., 2012) or LIF (Ai et al., 2020; Mi et al., 2001). Thus, interfering with EC migration, for example by exposure of the retina to VEGF signaling inhibitors (Morita et al., 2017), leads to a delay in astrocyte maturation. Conversely, mutations affecting astrocyte migration or maturation alter the formation of the VEGF gradient (by changing its topographical coordinates), thus resulting in an impaired EC chemoattraction (Arnold et al., 2012; Gnanaguru et al., 2013; Kautzman et al., 2018; Tao and Zhang, 2016).

The fact that lowered PDGFRα signaling leads to delayed EC migration, both in *Fat1*^*Nestin*^ retinas and in situations of biochemical or genetic inhibition of PDGFRα (Fruttiger et al., 1996; Tao and Zhang, 2016), suggests that VEGF production by immature astrocytes is also regulated by PDGFRα signaling. This supports a model in which there would be a threshold of PDGFRα signaling level for robust induction of VEGF expression (threshold above which VEGF can be induced/sustained), in addition to the oxygen/hypoxia threshold silencing VEGF expression in contact with ECs. Thus, a small change in the slope of PDGFRα level, such as the one observed in *Fat1*^*Nestin*^ retinas, may result in a positional change in the coordinates of the ON and OFF boundaries, and thus in the width of the VEGF^+^ zone, consequently affecting the slope of the VEGF gradient itself.

Another interesting phenotype is the presence of stochastic “avascular” patches, in which both astrocytes and endothelial cells are missing, suggesting a rupture of adhesion between astrocytes and the underlying layers. Such holes were also seen in context of mutations affecting the integrity of the inner limiting membrane (ILM). The ILM is a layer of extracellular matrix (ECM), enriched in laminins and collagens, delimiting the border between the inner surface of the retina and the vitreous cavity (Halfter et al., 2008). Mutations in several of these ECM components, such as *Lama1* (Edwards et al., 2010; Edwards et al., 2011) or *Lamb2/c3* (Gnanaguru et al., 2013), or in ECM receptors such as the dystroglycan complex (Clements et al., 2017; Tao and Zhang, 2016), alter ILM integrity, and are associated with vitreoretinopathies (Clements et al., 2017; Edwards et al., 2010; Edwards et al., 2011; Gnanaguru et al., 2013). The ILM provides a substrate for the migration of astrocyte progenitors, which navigate between the ILM and Müller glia, and mutations affecting ILM integrity also interfere with astrocyte migration, consequently perturbing retinal angiogenesis (Gnanaguru et al., 2013; Tao and Zhang, 2016). Whereas some of the ECM components are produced by the lens, others are secreted and modified by Müller glia, and the disruption of this process is sufficient to alter astrocyte migration and angiogenesis (Halfter et al., 2008; Tao and Zhang, 2016). Future studies will determine if altered ILM integrity can indeed be a potential cause for some of the a-vascular patches observed in *Fat1*^*Nestin*^ retinas and whether this phenotype results from suppression of *Fat1* activity in astrocytes or in Müller glia.

### Impact for vascular pathologies

Retinal vascular pathologies are becoming a major medical challenge, with the widespread occurrence of age-related macular degeneration, diabetic retinopathies or stroke (Quaegebeur et al., 2011). The present study identifies *Fat1* as a novel modulator of astrocyte maturation, the deletion of which in neural lineages impacts on the astrocyte/endothelial cell coupling, thus delaying postnatal angiogenesis, and leads to persistent vascular phenotypes in adult retinas. At present, we haven’t distinguished whether these adult phenotypes represent late consequences of the early abnormalities, or whether they highlight additional roles of *Fat1* during adult vasculature homeostasis. The increased vascular density and the presence of large collecting vessels in deep layers of *Fat1*^*Nestin*^ retinas are consistent with the possibility of neovascularization events occurring at adult stages, such as those occurring after transient ischemic events or vessel ruptures, owing to the capacity of hypoxia to induce VEGF expression. Although so far, none of the genome-wide association studies has identified *FAT1* mutations as being causally involved in any of the known human retinal pathologies, genetic and/or regulatory *FAT1* alterations have been associated with Facioscapulohumeral muscular dystrophy (FSHD), a neuromuscular pathology frequently associated with retinal vascular symptoms (Caruso et al., 2013; Mariot et al., 2015; Puppo et al., 2015). Results from the present study, by linking *Fat1* dysfunction to defective neurovascular communication, emphasize the key roles of astrocytes, Müller glia and neurons in the maintenance of vascular integrity. Furthermore, they uncover neural *Fat1* activity as a new component in the cross-talk between neural and endothelial cells, to ensure vascular integrity in the retina.

## Experimental procedures

### Mice

Animals were maintained and sacrificed in accordance with institutional guidelines. All procedures involving the use of animals were performed in accordance with the European Community Council Directive of 22 September 2010 on the protection of animals used for experimental purposes (2010/63/UE). The experimental protocols were carried out in compliance with institutional Ethics Committee guidelines for animal Research (comité d’éthique pour l’expérimentation animale–Comité d’éthique de Marseille) and in compliance with French law, under an agreement (Number D13-055-21) delivered by the “Préfecture de la Région Provence-Alpes-Côte-d’Azur et des Bouches-du-Rhône. Mice were sacrificed at P3, P5, or P7 for postnatal studies, whereas for collection of adult samples, they were first anesthetized with lethal doses of Ketamin and Xylasine, and euthanized by transcardiac perfusion with 4% PFA.

The mouse lines used in this study are described in supplementary Table S1, providing for each the Mouse Genome Informatics (MGI) reference number and link to the information page, a brief description of how it was made, and the reference to the article describing their production. These include three mutant alleles of the *Fat1* gene, all described and characterized in our previous studies (Caruso et al., 2013; Helmbacher, 2018): 1) A *Fat1*^*LacZ*^ genetrap allele (*Fat1*^*Gt(KST249)Byg*^, (Leighton et al., 2001)), in which *LacZ* expression reproduces that of the endogenous *Fat1* locus; 2) a *Fat1*^*Flox*^ conditional allele (*Fat1*^*tm1.1Fhel*^), in which exons 24 and 25 (the latter containing the transmembrane domain) are flanked by LoxP sites; 3) the constitutive knockout allele *Fat1*^*-*^ (also known as *Fat1*^*ΔTM*^, or *Fat1*^*tm1.2Fhel*^) derived from the *Fat1*^*Flox*^ allele by permanent cre-mediated deletion. Tissue-specific ablation in the neural lineage was achieved with the *Nestin-cre* line (Tg(Nes-cre)1Kln line, (Tronche et al., 1999)). We used the *Rosa26-YFP* line (Gt(ROSA)26Sor^tm1(EYFP)^Cos line (Jackson laboratory mouse strain 006148, (Srinivas et al., 2001)) as reporter of cre-mediated activity. Some of the embryos in which we study the coloboma phenotype also carry the transgenic line *MLC3F-2E-LacZ* (also known as Tg(Myl1-lacZ)1Ibdml, (Kelly et al., 1995; Kelly et al., 1997)). Genotyping is carried out as previously described for each line, except for new genotyping oligonucleotides allowing specific detection of the *Nestin-cre* transgene: Nes-Fw: 5’ AAG GGT TAA GGC CTA GGG ACC GCC C 3’; Nes-Rev (beginning of cre, reverse orientation): 5’ TTC AGG TTC TGC GGG AAA CCA TTT CC 3’.

As recently mentioned in a survey of various cre-lines (Luo et al., 2020), the *Nestin-cre* line exhibits a noteworthy amount of activity not only in the female germline (as previously known (Haigh et al., 2003)), but also in the male germline. Thus, when the transgene is carried by the same individual as the Floxed allele, Cre activity in the male germline can lead to the transmission by males of a *Fat1*^*-*^ recombined allele instead of the non-recombined allele (approximately 50% in the present study). Presence or absence of the recombined allele was evaluated by PCR. The crosses meant to produce mutants and littermate controls involved a *Nestin-cre; Fat1*^*Flox/*+^ or *Nestin-cre; Fat1*^*Flox/Flox*^ male, mated with *Fat1*^*Flox/Flox*^ females, which therefore represented a source of non-recombined allele. Thus, mutants derived from these crosses can be *Nestin-cre; Fat1*^*Flox/Flox*^ or *Nestin-cre; Fat1*^*Flox/-*^ (the two genotypes being pooled in our quantifications, and collectively called *Fat1*^*Nestin*^), whereas controls can be *Fat1*^*Flox/Flox*^, *Fat1*^*Flox/-*^ (the two genotypes being also pooled), even when they did not inherit the cre transgene. We also included some *Nestin-cre; Fat1*^*Flox/*+^ or *Nestin-cre; Fat1*^+*/*+^ controls in adult studies and obtained similar results.

### Tissue collection and immunohistochemistry (IHC)

Embryos were collected in cold PBS (on ice), and fixed in 4% paraformaldehyde (PFA) (in Phosphate buffered saline (PBS)) for 3-5 hours. Retinas from neonate mice were dissected from eyes previously immersed for 1 hour in 4% PFA, and were post-fixed for 1-2 hours maximum. Retinas from adult mice were dissected after transcardiac perfusion with fresh 4% PFA, and post-fixed for 1-2 hours as well. All tissues/embryos were then rinsed in PBS, and were either used directly for whole-mount immunohistochemistry (IHC) (free floating retinas), or cryoprotected and embedded/frozen for cryostat sectioning to perform IHC on sections.

In case of preparations for cryosections, tissues were cryo-protected by overnight immersion in 25% sucrose in PBS at 4°C, and embedded in a mix of 7.5% Gelatin; 15% sucrose (maintained liquid in a 42°C bath, and solidified on ice after adding the tissue). Once solidified, the position/orientation of the sample was indicated with a marker, prior to freezing the sample in isopentane, kept cold in a carbo-ethanol bath (dry ice/ethanol). Frozen samples were cryosectionned with a Zeiss Cryostat (10um thick), and sections were kept at −20°C until use.

For IHC, cryostat sections or free floating retinas were first incubated in PBS, 0.3% triton-X-100 for permeabilization. They were then subjected to a bleaching step in 6% hydrogen peroxide (H202) (mixing 1 Volume of 30% H2O2 with 4 Volumes PBS, 0.3% triton), for 30 minutes. For some antibodies (listed in Supplementary Table 2), a step of heat induced epitope retrieval (HIER) of cryostat sections was carried out prior to antibody incubation, by incubation in HIER solution at 95°C for 5-10 minutes. The HIER solution (0.1M Citrate buffer pH6.0; 0.2% tween-20) was preheated until it started boiling. Sections, pre-stabilized in cold HIER solution, were placed in the boiling HIER solution for a fixed time. Sections or floating retinas were then rinsed again (3 times) in PBS containing 0.3% triton-X-100, and incubated in a blocking solution containing: 20% Newborn Calf Serum (NCS), 0.3% Triton X-100, and PBS, together with the desired antibodies. When performing staining with the Isolectin IB4, 1mM of CaCl2 and 1mM MgCl2 were added to all solutions, to allow efficient Lectin binding.

Incubations with primary antibodies or Lectin were carried out in blocking solution (with Ca/Mg), overnight at 4°C, under agitation. All antibodies and binding reagents, references and conditions of use are listed in Supplementary Table 2. Following incubation with primary antibodies, retinas were rinsed multiple times in PBS 0.3% Triton X-100 (at least 5 times 15 minutes). Incubations with secondary antibodies were performed in blocking solution for 1h30 at room temperature, under agitation (or overnight at 4°C), after which samples were rinsed again multiple times (min 5 times) in PBS 0.3% Triton X-100. After completing the immunostainings, Retinas were flat mounted and covered with Mounting medium containing Prolong Gold anti-fade reagent and DAPI (Table S2). Images were acquired on a Zeiss AxioImager Z1 equipped with Apotome.

### Imaging and Quantification methods

Measurements of the relative eye area and grove width were carried out on images of embryos acquired with a Leica Stereomicroscope, and exploited using Axiovision software, following the procedure described previously (Helmbacher, 2018). The wild type and *Fat1*^*-/-*^ embryos were the same sample set as those in which we previously analyzed muscle development, thus sharing trunk length normalizing values (Helmbacher, 2018).

Quantifications of signal intensities were carried out using an approach similar to the method we previously described for quantifying in situ hybridization signal (Fan et al., 2015; Helmbacher, 2018). 10x mosaic images of flat-mounted retinas encompassing a quadrant were acquired, using an identical exposure time for all analyzed samples (ex, 290ms for PDGFRα intensity analysis). Images were rotated, such that a region of interest (ROI), encompassing a radius from the retina center to the periphery, is positioned horizontally, with the center to the left. A fixed size ROI (300×3000 pixels, representing 200×2000μm) was cropped, positioning its left border at the inner extremity of the optic stalk. Signal intensity along the entire ROI was then analyzed using ImageJ (analyze>plot profile), and data were exported for statistical analysis. Raw values (intensity ranging from 0 to 250) were then processed as described (Fan et al., 2015; Helmbacher, 2018). Since all ROIs included a peripheral region with minimal astrocyte coverage, this was used to define a fixed window (coordinates 2500-3000) for baseline intensity measurement (the baseline being the median value in the window, representing the background intensity). The window was manually adjusted in case of obvious artefact (damaged or folded retinal border). A median baseline was calculated for all controls of a same experimental series, and this value was subtracted from intensity measurement of each sample. The resulted intensity was thresholded at 0 (to avoid negative values). The maximal value was measured for each sample, in a window excluding the central ring (the thickness/intensity of which depending on dissection). The maximal amplitude was calculated by subtracting the baseline from the maximal value. A median maximal amplitude (MMA) was calculated for controls (MMAc) in each experimental group, and used for normalizing all data. The intensity was thus expressed as percentage of the MMAc (by dividing the thresholded intensity by the MMAc), varying along the proximo-distal coordinates of the ROI. For each mouse, the sum of intensity (area under the curve) was then calculated, either for the full radius (pixels 350 to 3000, excluding the inner ring), or for windows of fixed width (pixels 350 to 700; 800 to 1400; 1500 to 2000, for PDGFRα; and 100-500; 500-1000 and 1000-1500 for GFAP).

For the angle distribution analysis, a set of ∼100 PAX2^+^ nuclei was analyzed per mouse at the migration front in the same ROI (from the retina center to the periphery, 300×3000 pixels) from mosaic images of whole retinas. For each mouse, we calculated the percentage of nuclei displaying an angle in each angle category (ranges: 0-20°; 20-40°; 40-60°; 60-80°; 80-100°; 100-120°; 120-140°; 140-160°; 160-180°, with 90° representing the main outward direction (radius), in the width of the considered ROI. Thus, in each angle range, the analysis allowed retrieving one output value per mouse. Scatter plots showing each value are shown for the 80-100° range, and for a large angle group cumulating 0-60° and 120-180°.

Measurement of vessel length in adult retinas was carried out on Maximal intensity projection images of individual retinal layers, by applying the NeuronJ plugin in ImageJ (normally designed to trace axons) (Popko et al., 2009). For images of fixed size, the total length was calculated by addition of all measured distances. Other measurements (number and type of intersections), were done manually.

### Statistics

All measurements above led to retrieving one numerical value per mouse or embryo (measured value, ratio or percentage). This allowed comparisons between two groups (genotypes). For these comparisons, statistical significance was evaluated using either the unpaired Student t test, when data were showing a normal distribution and equal variance, or using the non-parametric Mann Whitney test otherwise. Differences were considered significant when p < 0.05. All p-values are indicated in the figures, except in Fig. 7C where ** indicates p-value < 0.001. For signal intensity curves (Fig. 6C), median values ± standard deviations are shown (in light blue, or light red). For scatter plots, all individual data are plotted (one per mouse), and the median value is shown with a black bar.

## Acknowledgements

I thank Dominique Fragano for mouse colony genotyping, the IBDM animal facility for mouse husbandry, past members of the Helmbacher lab for discussions, Flavio Maina and Robert Kelly for expert reading of the manuscript. Imaging was performed on PiCSL-FBI core facility (IBDM, AMU-Marseille) supported by the French National Research Agency through the “Investments for the Future” program (France-BioImaging, ANR-10-INBS-04).

## Supplementary Information

### Supplementary Figures

**Fig. S1.**
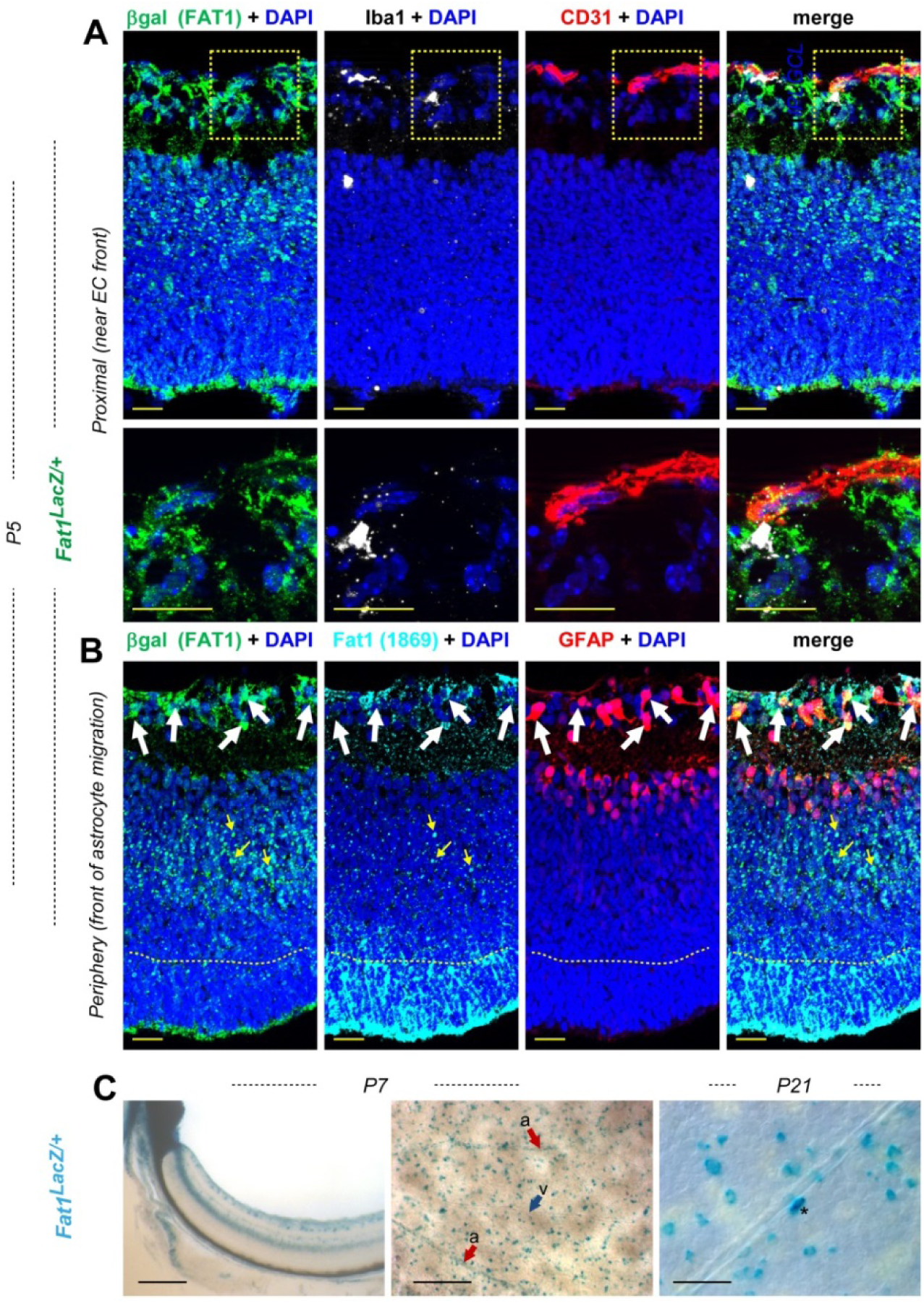
Complementary analysis of *Fat1* expression using the *Fat1*^*LacZ*^ reporter allele in retinas from P5, P7, and P21 mice. (A, B) Analysis by IHC on cross-sections of P5 *Fat1*^*LacZ*^ retinas was carried on with antibodies against β-galactosidase (green, A, B), CD31 (red, A), IBA1 (white, A), PDGFRα (red, B) and FAT1 (Cyan, B). Cell nuclei were labeled with DAPI. The bottom row panels in (A) show high magnification of the regions highlighted with yellow-dotted areas, focusing on endothelial cells and microglia, exhibiting low FAT1/ β-galactosidase levels, relative to nearby astrocytes. In (B), the apparent discrepancy between β-galactosidase and FAT1 localisation likely reflects the fact that the mutant fusion protein (detected by both antibodies) is retained in the secretory pathway, whereas only the wild-type protein reaches the membrane, thus decorating apical ends of Mueller glia, as well as light-receiving segments of rods and cones. (C) β-galactosidase activity is detected in P7 and P27 *Fat1*^*LacZ*^ retinas by X-gal staining. Flat mounted presentations allow detecting signal in scattered cells throughout the RGC layer, as well as along arteries (a) but not veins (v). Scale bars: (A, B): 20μm (C) P7: 200μm; P21: 50μm.

**Fig. S2.**
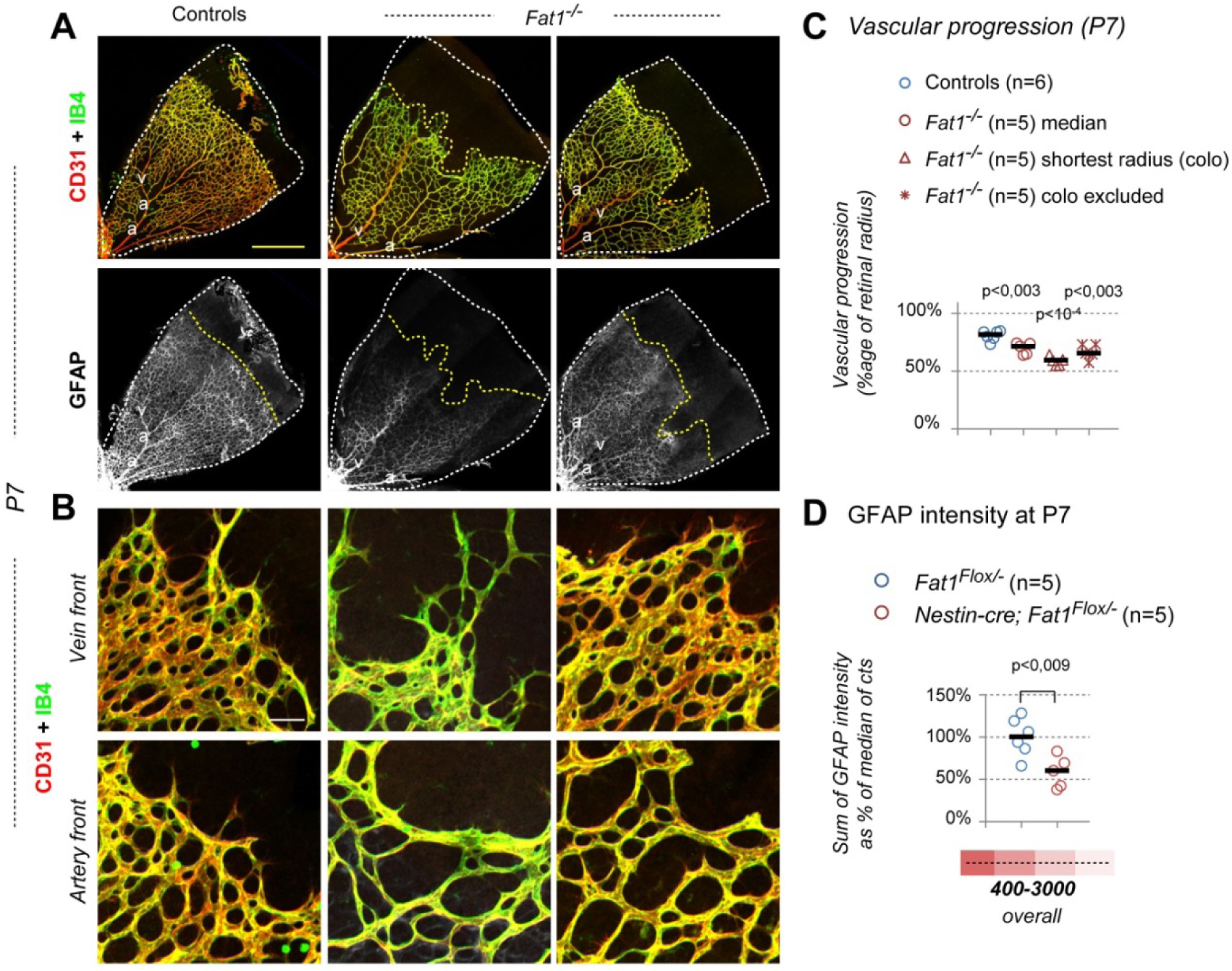
Constitutive Fat1 disruption impairs vascular progression p. (A, B) Flat mounted retinas from Wild type control, and two examples of *Fat1*^*-/-*^ mutant pups at P7, after immunohistochemistry with the GS-IB4 lectin (green), and antibodies against CD31 (red), and GFAP (white). Images in (A) feature an entire quadrant of retina; Lower pictures in (B) correspond high magnification views of the vascular front, in areas perfused by vessels connected to the nearest vein (top) or artery (bottom), respectively, showing the difference in EC density. (C) quantifications of the distance covered by IB4-positive ECs along the center-to-periphery axis, relative to the retinal radius at P7, distinguishing quadrants with minimal radius (with coloboma), from normally sized quadrants. (D) The intensity of GFAP staining was quantified along the radius was measured in control and *Nestin-cre; Fat1*^*Flox/-*^ mutants retinas at P7, and the sum of signal intensity was calculated for a window spanning the distances schematically represented corresponding to images shown in Fig. 4B. Scale bars: (A): 500μm (B) 50μm.

**Table S1.**
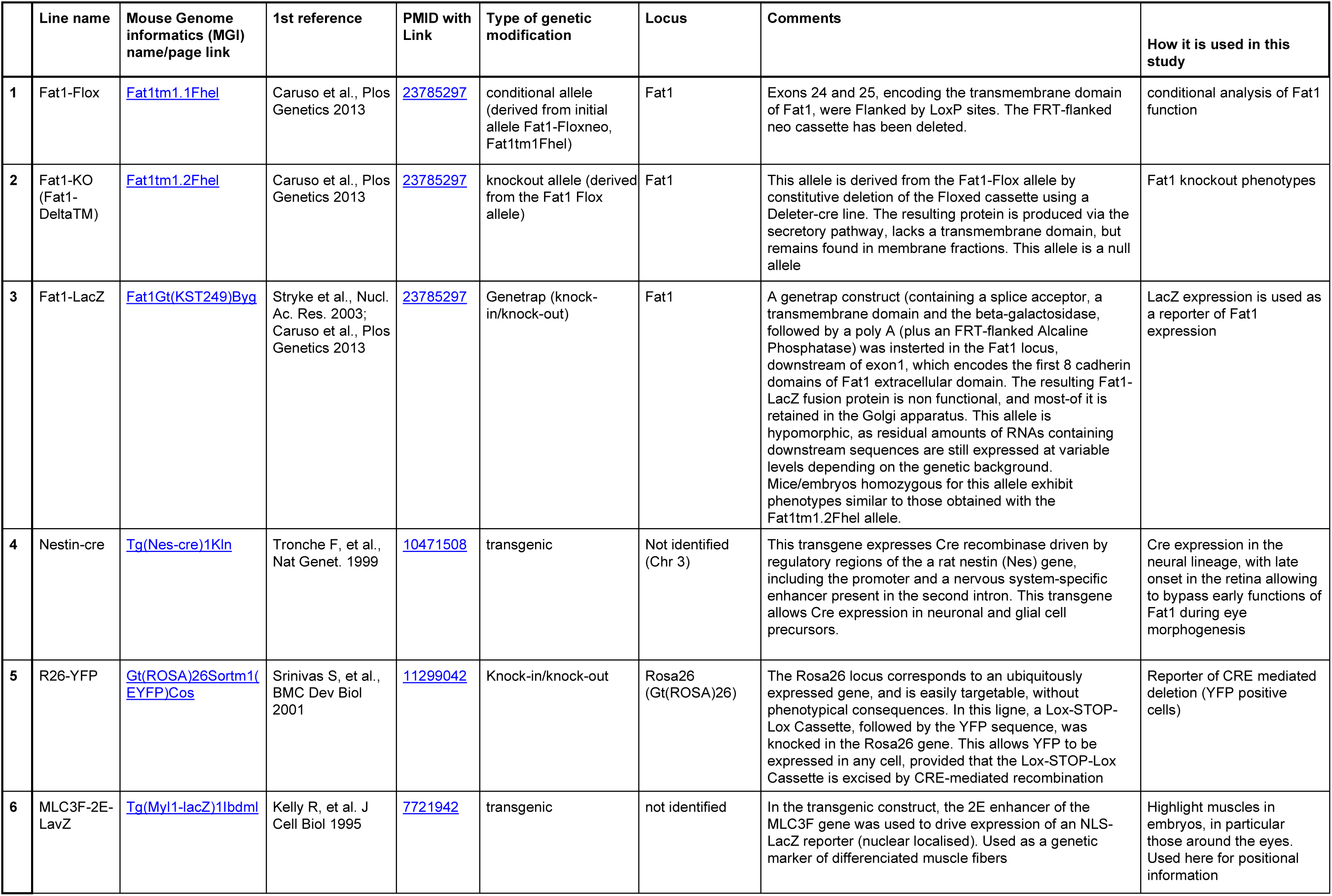
Mouse lines.

**Table S2:** Antibodies and other reagent.

| **Antibodies** | **provider** | **catalog number** | **Identifier (RRID)** |
| --- | --- | --- | --- |
| Anti-PAX7 (supernatant) Mouse Monoclonal antibody | DSHB | PAX7 (supernatant) | AB 2299243 |
| Anti-Neurofilament Mouse Monoclonal antibody, Clone 2H3 (bioreactor) | DSHB | 2H3 (bioreactor) | AB 531793 |
| Anti-Neurofilament Rabbit polyclonal Antibody | Millipore | Ab1987 | AB 91201 |
| Anti myosin heavy chain, type I (Myh1) Antibody, Mouse monoclonal, Clone MF20 (bioreactor) | DSHB | MF20 (bioreactor) | AB 2147781 |
| Anti-β-galactosidase Rabbit polyclonal antibody | Abcam | ab9361 | AB 307210 |
| Anti-GFP Antibody, Rabbit Polyclonal | Aves | GFP-1020 | AB 10000240 |
| Mouse Anti-Human GLUL (Glutamine Synthetase (GS)) Monoclonal Antibody, Unconjugated, Clone 2B12 | Novus | H00002752-M01A | AB 1199735 |
| Mouse Anti-Glial Fibrillary Acidic Protein (GFAP) Monoclonal Antibody, Unconjugated, Clone G-A-5 | SIGMA ALDRICH | G3893 | AB 477010 |
| Purified anti-Pax2 Rabbit polyclonal antibody | Covance | PRB-276P | AB 291611 |
| Purified Rat anti-mouse CD140a (Pdgfra) monoclonal antibody - clone APA5 | BD Biosciences | 558774 | AB 397117 |
| Rat anti-Mouse CD31 (PECAM) monoclonal antibody, Clone MEC13,3 (15.625 µg/ml) | BD Biosciences | 550274 | AB 393571 |
| Rat anti-Mouse CD31 (PECAM) monoclonal antibody, Clone MEC13,3 (0.5 mg/ml) | BD Biosciences | 550274 | AB 394816 |
| Anti-alpha, smooth muscle actin (Acta2), Rabbit monoclonal antibody (clone D4K9N) | Cell Signaling Technology | 19245S | AB 2734735 |
| Anti-Claudin 5 (CLDN5) Monoclonal Antibody (Clone 4C3C2), Alexa Fluor 488 conjugated | Thermofisher Scientific | 352588 | AB 2532189 |
| Rat Anti-PLVAP (Panendothelial Cell Antigen) Monoclonal Antibody, Unconjugated, Clone MECA-32 | BD Biosciences | 553849 | AB 395086 |
| Anti-IBA1 Rabbit polyclonal Antibody | WAKO CHEMICALS | 019-19741 | AB 839504 |
| **Other reagents** | **provider** | **catalog number** | **RRID** |
| Isolectin GS-IB<sub>4</sub> From *Griffonia simplicifolia*, Alexa Fluor™ 488 Conjugate | Thermofisher Scientific | I21411 | AB 2314662 |
| Prolong-Gold anti-fade Mountant, with DAPI (5x2ml) | Thermofisher Scientific | P36935 | - |

